# Environmental gradients reveal stress hubs predating plant terrestrialization

**DOI:** 10.1101/2022.10.17.512551

**Authors:** Armin Dadras, Janine M. R. Fürst-Jansen, Tatyana Darienko, Denis Krone, Patricia Scholz, Tim P. Rieseberg, Iker Irisarri, Rasmus Steinkamp, Maike Hansen, Henrik Buschmann, Oliver Valerius, Gerhard H. Braus, Ute Hoecker, Marek Mutwil, Till Ischebeck, Sophie de Vries, Maike Lorenz, Jan de Vries

## Abstract

Plant terrestrialization brought forth the land plants (embryophytes). Embryophytes account for most of the biomass on land and evolved from streptophyte algae in a singular event. Recent advances have unraveled the first full genomes of the closest algal relatives of land plants; among the first such species was *Mesotaenium endlicherianum*. Here, we used fine-combed RNAseq in tandem with photophysiological assessment on *Mesotaenium* exposed to a continuous range of temperature and light cues. Our data establish a grid of 42 different conditions, resulting in 128 transcriptomes and ~1.5 Tbp (~9.9 billion reads) of data to study combinatory effects of stress response using clustering along gradients. We describe major hubs in genetic networks underpinning stress response and acclimation in the molecular physiology of *Mesotaenium*. Our data suggest that lipid droplet formation, plastid and cell wall-derived signals denominate molecular programs since more than 600 million years of streptophyte evolution—before plants made their first steps on land.

## MAIN

Plant terrestrialization changed the face of our planet. It gave rise to land plants (Embryophyta), the major constituents of Earth’s biomass (Bar-On et al. 2018) and founders of the current levels of atmospheric oxygen (Lenton et al. 2016). Land plants belong to the Streptophyta, a monophyletic group that includes the paraphyletic freshwater and terrestrial streptophyte algae and the monophyletic land plants. Meticulous phylogenomic efforts have established the relationships of land plants to their algal relatives (Wickett et al. 2014; One Thousand Plant Transcriptomes Initiative, 2019). These data brought a surprise: the filamentous and unicellular Zygnematophyceae—and not other morphologically more elaborate algae—are the closest algal relatives of land plants. Now the first three genomes of major orders of Zygnematophyceae (see Hess et al., 2022) are at hand: *Mesotaenium endlicherianum, Spirogloea muscicola* (Cheng et al., 2019), and *Penium margaritaceum* (Jiao et al., 2020). Using these, we are beginning to redefine the molecular chassis shared by land plants and their closest algal relatives. Included in this shared chassis will be those genes that facilitated plant terrestrialization. We here focus on one critical aspect: the molecular toolkit for the response to environmental challenges. For this, we use the unicellular freshwater/subaerial alga *Mesotaenium endlicherianum*.

Land plants use a multilayered system for the adequate response to environmental cues. This involves sensing, signaling, and response mainly by the production of, e.g., protective compounds. Some of the most versatile patterns in land plant genome evolution concerns genes for environmental adaptation (Golicz et al., 2016; Gordon et al., 2017; Bayer et al., 2020). That said, there is a shared core of key regulatory and response factors that are at the heart of plant physiology. These include phytohormones such as abscisic acid (ABA) found in non-vascular and vascular plants (for an overview, see Umezawa et al., 2010; Bowman et al., 2019), protective compounds resting on specialized metabolic routes such phenylpropanoid-derived compounds as well as proteins such as LATE EMBRYOGENESIS ABUNDANT (LEA; Hundertmark and Hincha 2008; Carella et al., 2019). Many of the genes integrated into these stress-relevant metabolic routes have homologs in streptophyte algae (Rieseberg et al., 2022). Taking angiosperms as reference, such stress-relevant pathways are often patchy. Whether these are also used under the relevant conditions is currently unknown. For example, while Zygnematopyhceae have a homolog to the ABA-receptor PYL (de Vries et al. 2018, Cheng et al., 2019), this homolog works in a different, ABA-independent fashion (Sun et al., 2019). Thus, it is important to put the genetic chassis that could act under environmental shifts to the test.

Here, we used a fine grid of a bifactorial gradient for two key terrestrial stressors, variation in irradiance and temperature, to probe the genetic network that the closest algal relatives of land plants possess for its responsiveness to abiotic environment. Correlating growth, physiology, and global differential gene expression patterns from 128 transcriptomes (9,892,511,114 of reads, 1.5 Tbp of data) across 126 distinct samples covering a temperature range of >20°C and light range of >500 μmol photons m^−2^ s^−1^, we pinpoint hubs in the circuits that have been shared along more than 600 million years of streptophyte evolution.

## RESULTS

### A physiological grid: co-dependency of eurythermy and euryphoty in *Mesotaenium*

We studied the genome-sequenced strain SAG 12.97 of the freshwater alga *Mesotaenium endlicherianum*, a member of the Zygnematophyceae, the closest algal relatives of land plants (Cheng et al., 2019; Figure 1a and 1b). We cultivated *Mesotaenium* in a large-scale setup in 1.5 liters of C medium up to a cell density of 0.33 AU at 680 nm. The culture was distributed across 504 wells (42 twelve well plates; 2.5 mL of culture per well). The well plates were placed on a table with a temperature gradient from 8.6±0.5 °C to 29.2±0.5 °C on the x-axis. On top of the table, white LED lamps created an irradiance gradient from 21.0±2.0 to 527.9±14.0 μmol photons m^−2^ s^−1^ across the y-axis, thus creating a 2D gradient table (Figure 1b, Suppl. Table 1). The 504 cultures were exposed to this gradient setup for 65 hours. The physiological status of the algae was assessed by determining the maximum quantum yield (*F*_*v*_/*F*_*m*_) using pulse amplitude modulation fluorometry (PAM; IMAGING PAM, Walz, Germany); growth was assessed using a microplate reader with absorption at 480 nm, 680 nm, and 750 nm (Figure 1c); the entire procedure was repeated in three successive biological replicates (i.e. three runs of the table, 504 *F*_*v*_/*F*_*m*_ and 4,536 absorption measurements per replicate).

**Figure 1:**
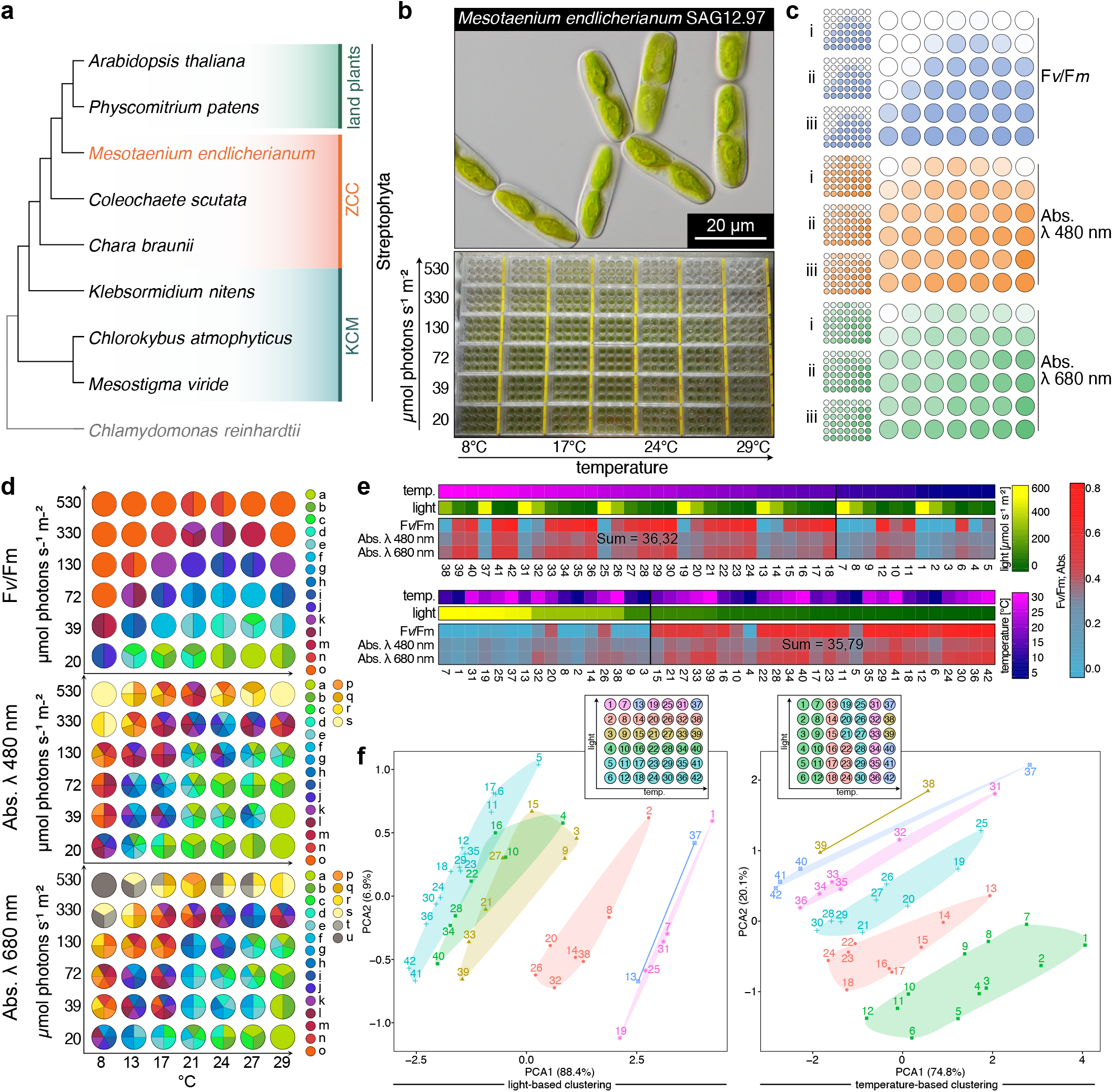
A fine-combed setup for assessing environmental cues in Mesotaenium. (a) Cladogram of Streptophyta, highlighting that *Mesotaenium endlicherianum* SAG 12.97 is a representative of the closest algal relatives of land plants. (b) *Mesotaenium endlicherianum* grown in C-medium in 42 twelve-well plates on a gradient table that produces a temperature range of 8.6±0.5 °C to 29.2±0.5 °C on the x-axis and an irradiance gradient of 21.0±2.0 to 527.9±14.0 μmol photons m^−2^ s^−1^ on the y-axis. (c) Overview of the measured maximum quantum yield *F*_*v*_*/F*_*m*_ as a proxy for gross physiology (blue) and Absorption (Abs.) at 480 (orange) and 680 nm (green); individual replicates of the biological triplicates are shown on the left and the average values are shown on the right. (d) Statistical analysis of the physiological values (*F*_*v*_*/F*_*m*_, Abs. 480 nm, Abs. 680 nm). Numbers correspond to environmental conditions on the table. Biological triplicates were grouped into significant groups (a-u) with R (version 4.1.3) using a Kruskal-Wallis test coupled with Fisher’s least significance; *p* values were Bonferroni corrected. Significant differences at *p* ≤ 0.001 are shown as letters. (e) Heatmaps displaying averaged physiological values of the 42 conditions sorted either by (i) temperature or (ii) light. A cut-off was set (black vertical line) based on the distribution of the highest values, which were then summed up to determine a positive correlation with temperature or light conditions. (f) Two principal component analyses (PCA) showing the correlation of light conditions (left) or temperature conditions (right) to physiological values (*F*_*v*_*/F*_*m*_, Abs. 480, 680 nm). Clusters are shown in different colors, which are also visualized in an overview scheme of the gradient table at the top of the plots.

The algae showed significant differences (p ≤ 0.001) in growth and gross physiology: *F*_*v*_/*F*_*m*_ values as well as absorption values decrease (for *F*_*v*_/*F*_*m*_ values at 20.5±1.0 °C: from 0.66±0.02 for I=21.14 μmol photons m^−2^ s^−1^ to 0.042±0.04 for I=534.7 μmol photons m^−2^ s^−1^) with rising intensities of irradiance (Figure 1d, Suppl. Fig. 1, Suppl. Table 2). The lowest *F*_*v*_/*F*_*m*_ values (down to zero) were recorded at conditions of highest irradiance and lowest temperature. Here, low temperature had a stronger negative impact on growth and physiology than light (for *F*_*v*_*/F*_*m*_ values: at 8.6±0.5 °C, 0.011±0.02 at 133±27 μmol photons m^−2^ s^−1^ compared to 0.463±0.02 at 29.2±0.5 °C at 118±25 μmol photons m^−2^ s^−1^). Values on growth and physiology clustered by light were less broadly distributed than if clustered by temperature (Figure 1e, 1f). Even the highest light intensity (527.9±14.0 μmol photons m^−2^ s^−1^) was stressful, but tolerable for the physiology of *Mesotaenium* at temperatures between 20.5±0.1°C (*F*_*v*_*/F*_*m*_=0.042±0.04) to 25.3±0.1°C (*F*_*v*_*/F*_*m*_= 0.045±0.04); more extreme temperatures resulted in undetectable Fv/Fm values. Thus, eurythermy might establish the foundation for euryphoty in *Mesotaenium endlicherianum*.

### Fine-combed global differential gene expression profiles and gene models for *Mesotaenium*

To shed light on the molecular mechanisms that underpin the switch from tolerable steady-state conditions to adverse environmental cues in *Mesotaenium*, we applied global gene expression analyses using RNAseq. We pooled all twelve wells per plate and extracted RNA from a total of 126 samples (42 plates, three biological replicates). 114 samples yielded usable RNA that was used to build 128 libraries (with a minimum of three biological replicates and additional technical replicates) for sequencing on the Illumina NovoSeq6000 platform. We generated a total of 1.5 Tbp of 150 bp paired read data at an average depth of 37.7 million reads per sample (~9.9 billion reads in total). Building on this wealth of data, we updated the *Mesotaenium* gene models. The number of protein-coding mRNAs increased from 11,080 in the original annotation (V1; Cheng et al. 2019) to 40,326 protein-coding mRNA (26,009 high confidence, 14,317 low confidence; including splice variants) in 19,233 genes; an additional 4,408 mRNA (in 4,312 genes) labeled as “predicted gene” in our gene models (Suppl. Table 3). The new gene models of annotation V2 brings the number of genes in *Mesotaenium* closer to other Zygnematophyceae with similar genome sizes; V2 has 43 more BUSCO genes (+10%; 21 less fragmented, 22 less missing; viridiplantae_odb10) than V1 (Suppl. Fig. 2). Besides, we calculated Annotation Edit Distance metrics (AED) to assess the congruence (0 to 1, with 0 being the best) between biological evidence and V1 and V2. In the cumulative fraction of annotation against AED score, V2 has more mRNAs with AED < 0.5. For example, 70% of mRNAs in V1 (7,756 mRNAs) have an AED score < 0.5 compared to 60% in V2 (26,840 mRNAs). This is sensible since V2 was built based on the same set of evidence used to calculate AED and it shows higher congruence with them (Suppl. Fig. 3). Thus, we pseudoaligned our data onto the new *Mesotaenium* transcriptome V2 (average alignment rate was 87.31%; Suppl. Table 4).

To understand the gross profile of the gene expression data, we performed principal component analysis (PCA; Figure 2a). Independent biological replicates from the same condition clustered in close proximity. The most variation in data was explained by temperature (PC1; describes 35% of variance), followed by irradiance (PC2; describes 18.1% of variance). We evaluated the distance (Figure 2b) and Spearman correlation (Figure 2c) using all genes to look for trends among different growth conditions. The data can be grouped into at least three categories: (1) samples with high light and/or high temperature, (2) a collection of low-temperature (8, 13, 17 °C) samples, and (3) samples at stead-state. Large clusters included steady-state, high light + heat, and high light. Most distinct was the cluster formed by samples from the high temperature + high light (Small multiples; Figure 2d and 2e).

**Figure 2:**
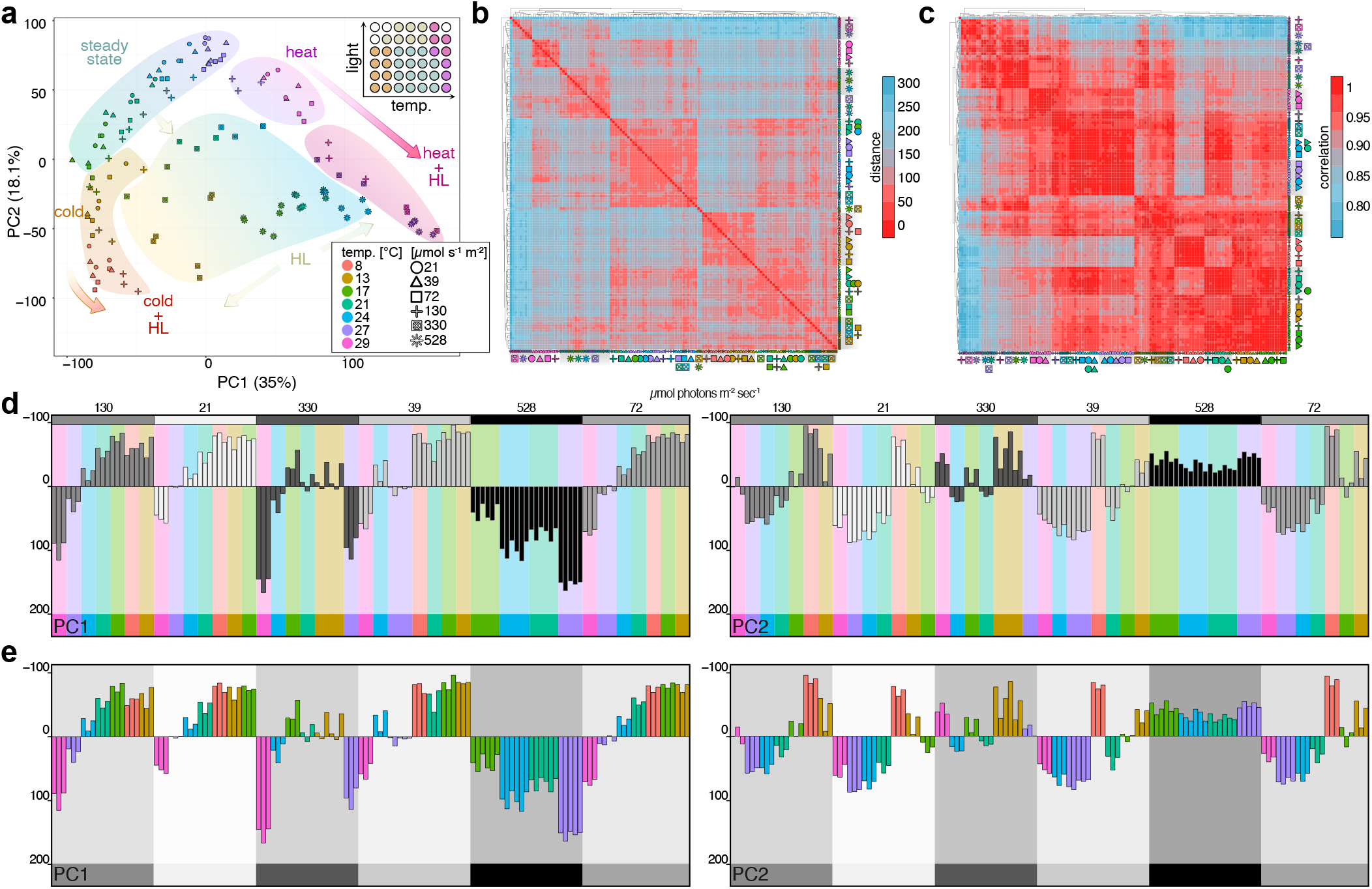
Global profiles of environment-governed gene expression response. (a) Principal component analysis (PCA), visualizing PC1 and PC2. Backgrounds were drawn to highlight our interpretation of the observed trends; samples are coded by color (temperature) and symbols (irradiance in μmol photons m^−2^ s^−1^). (b) Visualization of Euclidean distances between samples via heatmap, from red, zero distance, to blue, furthest distance (a distance of 300). (c) Heatmap of Spearman correlation between samples, from red, maximum correlation (1.0), to blue, least correlation (< 0.8). The clusters were calculated via the Euclidean distance. (d) PC1 and PC2 scrutinized using a small multiples method of light intensity and (e) temperature. In (d) shades of gray corresponds to different light intensities. In (e) different colors represent different temperatures and were mapped with the same colors as (a).

### Plastid-related genes stand out in differential gene expression profiles

For dissecting the differential gene expression responses, we divided the table into nine sectors and, additionally, a cohort of stressed algae based on *F*_*v*_*/F*_*m*_ < 0.5 (Figure 3). 36 comparisons were performed, among which we focused on nine, which additionally included the *F*_*v*_*/F*_*m*_-based comparison. Genes were considered to be differentially expressed between groups at an absolute fold change ≥ 2 and a Benjamini-Hochberg corrected *p* ≤ 0.01 (Figure 3a and b). Gross gene expression profiles were titratable by the intensity of environmental cues, i.e., with increasing disparity between conditions compared, and overall following the pattern in the PCA (*cf*. Figure 3b and Figure 2a). The most differentially regulated genes (6,578) were pinpointed by comparing low light and low temperature (LLI_LT) versus high light and high temperature (HLI_HT). Enriched GO terms among regulated genes most frequently included plastid biology-associated genes (Figure 3c). To scrutinize these data for specific genes that show a robust and universal response to alterations to the environment, we intersected all 8,157 significantly regulated genes pinpointed by the different comparisons. 3, 30, and 124 genes overlapped among all 9, 8, and 7 comparisons, respectively. These concertedly pinpointed genes were mostly light harvesting genes, corroborating the importance of plastids in the overall cell biology of *Mesotaenium* (Figure 3d). Indeed, the 30 genes found in all comparisons included for example reactive oxygen species (ROS)-relevant genes such as ELIP and fatty acid metabolic genes. To understand whether these genes integrate into the context of molecular programs, we next looked at gene co-expression.

**Figure 3:**
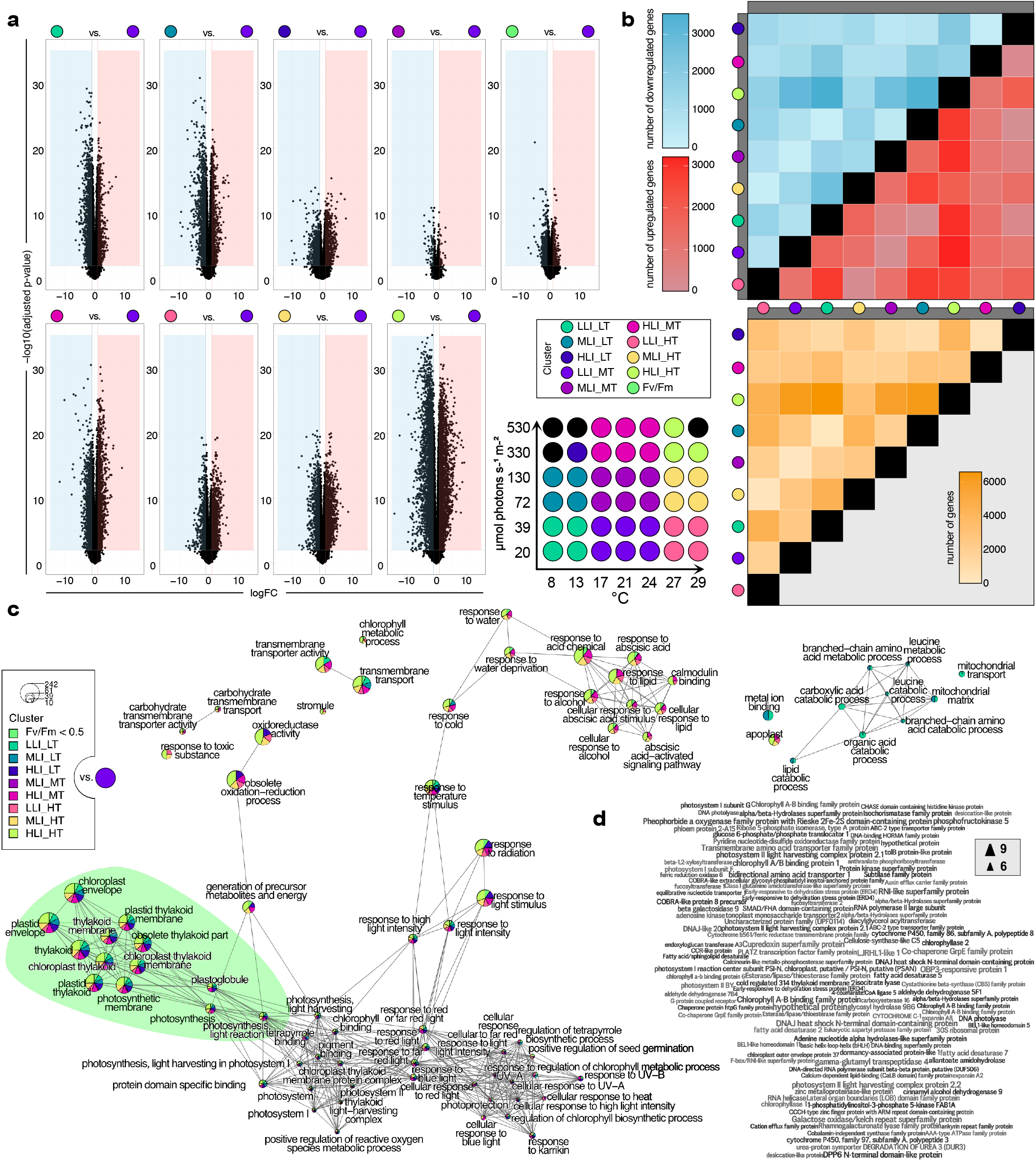
Stress-titratable global differential gene expression profiles. To perform differential gene expression analysis, we divided the table into 9 sectors (see scheme of the table); additionally, a tenth group was raised based on Fv/Fm < 0.5. Linear models were fitted for each gene and empirical Bayes statistics computed for differentially expressed genes (DEGs) by the limma package. In total, 37 comparisons were made. DEGs were defined as genes with an absolute fold change ≥ 2 and BH-adjusted *p* value lesser than 0.01. (a) Volcano plots of DEGs for 9 selected comparisons based on the sectors and the Fv/Fm < 0.5 criterion. (b) Heatmaps of numbers of DEGs for all sector-based comparisons (blue, downregulation; red, upregulation; yellow, sum of up- and down-regulated genes); grey bars label the first component (treatment) for calculating the contrasts (treatment vs. control). (c) Biological theme comparison summarizing all GO-term enrichment analysis with adjusted p-value ≤ 0.01 of DEGs against all genes that were expressed and passed the filtering in our analyses as background. The size of each circle is proportional to the count of each GO-term. Only the top 30 enriched terms are shown. (d) Wordle of the 124 genes that showed significant regulation across multiple comparisons shown in Figure 3a; word size correspond to the number of comparisons (based on (a)) in which a gene appeared.

### Unsupervised gene expression clusters recover genetic programs shaped by physiology

The environmental gradients triggered changes in the expression of gene cohorts. We wanted to understand their concerted action independent of any prioritization guided by homology to any land plant genes—solely from the molecular programs that operated in the algae. To do so, we applied weighted gene co-expression network analysis (WGCNA) for unsupervised clustering (Figure 4). To then understand the driving forces behind these changes, we turned to the highly connected genes (nodes) in the network—the hubs (Figure 5).

**Figure 4:**
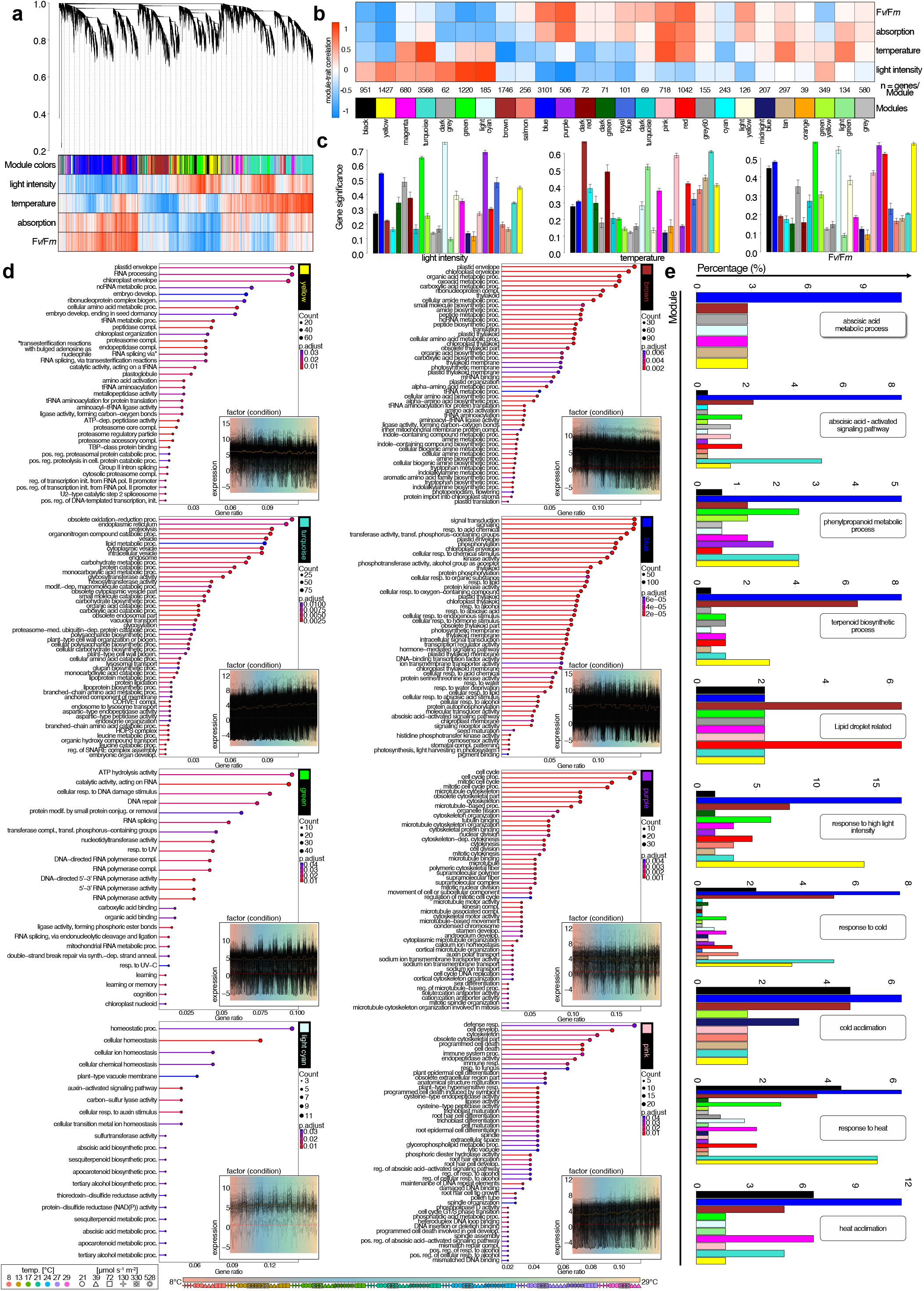
Unsupervised gene expression clusters recover genetic programs separated by environmental cues. Gene expression clustering into 26 colored modules was performed using WGCNA; grey is the module of unclustered genes. (a) Hierarchical cluster tree of 17,095 genes. The heat map below the dendrogram shows the gene significance measure (from red, positive correlation, to white, no correlation, to blue, negative correlation) for the four different conditions / physiological parameters. (b) Heat map of the module–trait correlation based on eigengenes (from red, positive correlation, to white, no correlation, to blue, negative correlation); see Suppl. Fig. 7 (c) Bar plots of the mean gene significance across modules (given in the corresponding module color) towards the parameters light intensity, temperature, and *F*_*v*_*/F*_*m*_. (d) Enriched GO-terms for eight of the 26 modules; each inset shows the gene expression profiles of all genes in a given module. (e) Arabidopsis homologs for key processes were mined based on keywords; they were retrieved from a look-up table of BLASTp hits in a search of *Mesotaenium* V2 against *A. thaliana* representative protein sequences. Bar charts show the percentage of detected *Mesotaenium* homologs across the modules relative to the number of all Arabidopsis IDs assigned to the terms. No blast hit was not depicted. Abbreviations: proc. = process; reg. = regulation; biogen. = biogenesis; develop. = development; pos. = positive; neg. = negative; init. = initiation; GEP = Gene expression profile; med. = mediated; dep. = dependent; modif. = modification; conjug. = conjugation; anneal. = annealing; compl. = complex; synth. = synthesis; resp. = response; transf. = transferring.

**Figure 5:**
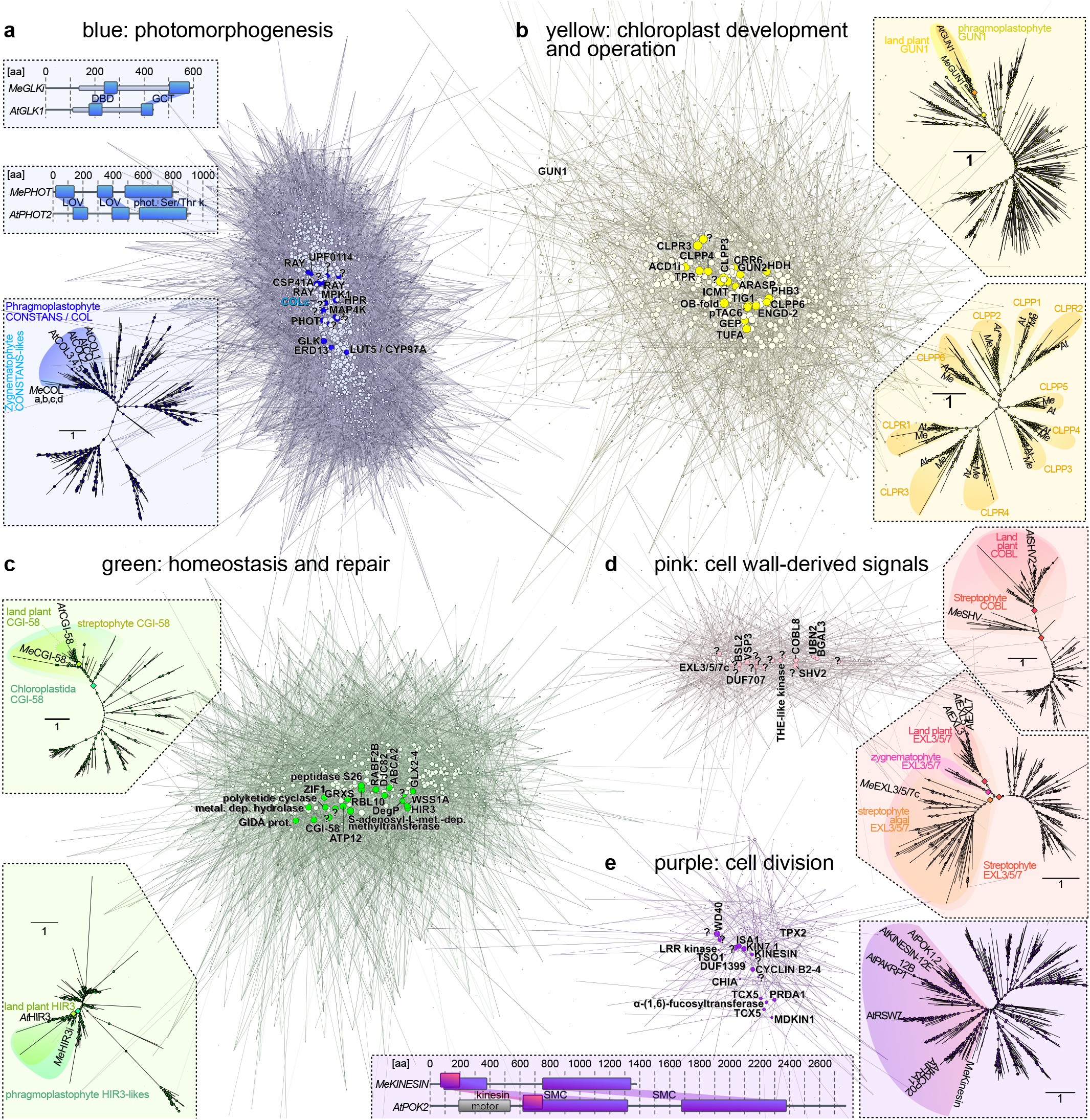
Molecular programs for environmental responses around recurrent plant hubs. Visualization of the co-expression networks clustered by WGCNA into the modules blue (3,101 genes), yellow (1,427 genes), green (1,220 genes), purple (506 genes), and pink (718 genes). Nodes, circles representing genes, are connected by edges whose weight is based on a weighted topological overlap matrix (TOM); weight is shown in a color gradient from light to dark indicating low to high topological overlap values in the TOM. Brightly colored nodes represent the 20 most connected genes (hubs) and are annotated; all other nodes are depicted in the corresponding paler color. Hubs are annotated based on homology. Around the clusters, different protein-coding hub genes are highlighted, giving information such as predicted domain structures or phylogenetic relationships; fully-labelled phylogenies are deposited in Supp. Fig. 26. Circles in phylogenies represent ultrafast bootstrap support, with larger circles represent high/full support; diamond symbols indicate high (>90) support for branches separating highlighted clades. An alignment of GLK homologs can be found in Suppl. Fig. 8.

The 17,905 genes expressed in our samples (and that passed the minimum expression threshold) were clustered into 26 modules, which we refer to with colors (Figure 4a). Orange is the smallest module (39 genes), the largest modules are Turquoise, Blue, and Brown with 3568, 3101, and 1746 genes, respectively. The samples were taken under a range of distinct physiological conditions. Resulting data are a combined expression of the different environmental cues and the modulation of the algal physiology. To investigate the biological role of each module, we used their eigengenes as representatives for the modules’ gene expression profiles and correlated their behavior with the two environmental cues light intensity and temperature as well as the algal parameters absorption (culture density and pigmentation) and *F*_*v*_*/F*_*m*_ (overall physiological status). One of the foremost general patterns in cellular response to stress are ROS. These act as signals as well as culprits that, if not quenched, damage biomolecules; this was represented in GO terms of module Green that positively correlated with light intensity (*r* = 0.88, *p* = 6×10^−43^) and negatively with *F*_*v*_*/F*_*m*_ (*r* = −0.79, *p* = 6×10^−29^) (Figure 4d and Suppl. Fig. 4 to 7 and Suppl. Table 5 and 6).

The clusters also recovered the genetic signatures of thriving algae. Module Purple negatively correlates with increasing light (*r* = −0.94, *p* = 3×10^−60^) and positively with absorption and *F*_*v*_*/F*_*m*_ (*r* = 0.67, *p* = 2×10^−18^ and *r* = 0.67, *p* = 2×10^−18^). These dense and physiologically healthy cell populations (experiencing no light stress) ramped up cell division (see Figure 4D and Suppl. Table 6), signified by homologs of cyclin and TPX2 appearing as hub genes. The 9th most connected hub gene was a kinesin homologous to important proteins such as PHRAGMOPLAST ORIENTING KINESIN 2 (Figure 5; Suppl. Table 7), which thus is a likely conserved cell division hub of all Phragmoplastophyta—going back to a common ancestor that lived in the late Cryogenium.

### Conserved hubs for plastid-derived signals

Chloroplasts act as environmental sensors in land plant cells (Kleine et al., 2021). In concert with this, many of the clusters we identified were associated with plastid biology and/or physiology (Figure 4d, Suppl. Fig. 4 to 7, Suppl. Table 6). The brown cluster showed many plastid-related terms and negatively correlates with temperature (*r* = −0.95, *p* = 7×10^−65^) (Suppl. Fig. 5) and showed enrichment in GO-terms related to plastids, general transcription and translation. Among the top 20 hub genes in cluster brown, 12 were associated with translation and ribosomes (Suppl. Table 7). The light cyan cluster positively correlates with increasing light (*r* = 0.93, *p* = 10^−56^) (Suppl. Fig. 6) and negatively with *F*_*v*_*/F*_*m*_ (*r* = −0.67, *p* = 5×10^−18^) (Suppl. Fig. 4) whereas the blue cluster negatively correlates with increasing light (*r* = −0.76, *p* = 10^−25^) and positively with *F*_*v*_*/F*_*m*_ (*r* = 0.67, *p* = 2×10^−18^). Concomitantly, the blue module had a high number of enriched GO-terms (Suppl. Fig. 5 and Suppl. Table 6), many of which were plastid-related terms, cellular signaling, and terms that tie the two together; that is, signaling processes emanating from the plastid. This was also prominent in the light cyan module, where several terms related to terpenoid and apocarotenoid metabolism were enriched.

The hubs of many clusters, including those blue, light cyan, and yellow mentioned before, reflect an association with plastid-related processes. To highlight a few, the second most connected gene in module Blue was a homolog of GLK1 (Suppl. Fig. 8), a transcriptional factor (TF) that regulates chloroplast development and the activity of nuclear genes involved in photosynthetic light reaction and chlorophyll biosynthesis (Rossini et al., 2001; Yasumura et al., 2005; Waters et al., 2009). Blue also featured hydroxypyruvate reductase, important in photorespiration (Timm et al., 2008), as the fourth most connected gene. A CYP450 gene homologous to LUTEIN DEFICIENT 5 (LUT5), was the 7^th^ most connected, suggesting the involvement of pigment-related signaling. Moreover, a homolog of ABA responsive elements-binding factor 2 (ABF2) was part of cluster Blue, bolstering previous discussions that parts of the ABA signaling module consist of ancient wires whose relevance in stress response predate plant terrestrialization and ABA dependency (de Vries et al., 2018; Sun et al., 2019; Fürst-Jansen et al., 2020).

Next to GLK—the most connected TF—other highly connected TFs appeared in Blue. These included the photomorphogenesis-regulating CONSTANS-like 3 (COL3; 4^th^ most connected TF). Noteworthily, also a homolog of CONSTITUTIVE PHOTOMORPHOGENIC 1 (COP1) was present in module Blue; CO/COL and GLKs are both degradation targets of COP1 (Liu et al., 2008; Sarid□Krebs et al., 2015; Ordoñez-Herrera et al., 2018). Further, the circadian regulator BROTHER OF LUX ARRHYTHMO (2^nd^ most connected TF). Further, homologs of ETHYLENE-INSENSITIVE3-like 1 (6^th^ most connected TF) and several ERFs were among the most connected TFs. A link to ethylene is noteworthy, because investigations of the Zygnematophyceae *Spirogyra pratensis* (*Sp*) have shown that *SpEIN3* can rescue Arabidopsis *ein3-1* mutant plants (Ju et al., 2015). Furthermore, exogenous application of ethylene on *Spirogyra* triggers stress-, plastid- and photosynthesis-associated gene expression responses similar to land plants (Van de Poel et al., 2016). This speaks to a conserved regulatory framework that involves ethylene-associated factors, and maybe ethylene itself, in environmental signaling cascades in the common ancestor of and plants and their closest algal relatives.

Light cyan featured not only hubs related to ROS homeostasis from the thioredoxin superfamily and other light-induced proteins, but also pigment and apocarotenoid metabolism; these are the source of important signals from the chloroplast that likely have deep evolutionary roots (Rieseberg et al., 2022) and are also formed by light dependent oxidative reactions (recently reviewed by Moreno et al., 2021). Module Yellow correlated positively with light intensity (*r* = 0.62, *p* = 10^−14^) and negatively with absorption and *F*_*v*_*/F*_*m*_ (*r* = −0.79, *p* = 10^−28^ and *r* = −0.81, *p* = 3×10^−31^; Figure 3B); GO terms associated with plastids and proteolytic enzymes (FtsH, ClpP; Kato et al., 2012), recapitulating well-known ties of protein homeostasis and plastid maintenance. Indeed, cluster yellow featured five hubs that are homologous to CLP proteases, critical for chloroplast protein homeostasis (Sjögren et al., 2006; Nishimura et al., 2016), and hubs homologous to genes that orchestrate the coordination of transcriptional activity between chloroplasts and the nucleus; the latter included homologs of (i) pTAC6, which is essential for plastid gene expression and thus chloroplast development in Arabidopsis (Pfalz et al., 2006), and (ii) a homolog of GENOMES UNCOUPLED 2, one of the foremost genes in the classical plastid–nucleus communication pathway (Susek et al., 1993). Among the TFs in cluster yellow was a homolog of the bZIP light signaling master regulator ELONGATED HYPOCOTYL 5 (HY5; reviewed in Jiao et al., 2007).

### Of ancient signaling cascades and cell wall perturbance

Mitogen-activated protein kinases (MAPK) constitute environmental response pathways in all eukaryotes (Chen and Thorner, 2007). In land plants, several abiotic and biotic cues have been described to trigger MAPK-mediated signaling (Nakagami et al., 2005; Rodriguez et al., 2010; Meng and Zhang, 2013; Chen et al., 2021); MAPK and phototropin kinases appeared as hubs in cluster Blue. Moreover, plant MAPK-based signaling is interwoven with wound response and brassinosteroid signaling (Nakagami et al., 2005). Stress often coincides with a perturbance of plant cell wall homeostasis. Cluster Pink includes hubs for such wounding and cell-wall derived signals. This was paired with the GO term brassinosteroid signaling, which balances growth, cell wall homeostasis, and stress in *Arabidopsis* (Sun et al., 2010; Planas-Riverola et al., 2019). Among the hubs in cluster Pink were homologs for (i) diverse receptor kinases known from *Arabidopsis* to sense alterations in cell wall integrity (Hématy et al., 2007), and (ii) EXORDIUM (of which *Mesotaenium* has 12 homologs), which integrates growth with environmental signaling (Schröder et al., 2009). This was paired with the COBRA family proteins being the most and third most connected hubs in the module. These proteins are known to be involved in cell expansion and balancing pathogen response with growth (Schindelmann et al., 2001; Roudier et al., 2002; Ko et al., 2006). It appears that *Mesotaenium* bears parts of a loop that senses physico-chemcial perturbance of cell wall homeostasis; in land plants, these loops include brassinosteroid signaling (Wolf et al., 2014).

### Lipid droplet formation constitutes a stress response predating plant terrestrialization

In land plants lipid droplet (LD) formation and triacylglycerol (TAG) accumulation is common to many stress responses, including heat, cold and drought (Higashi et al., 2015; Mueller et al., 2015; Gidda et al., 2016; Doner et al., 2021; Krawczyk et al., 2022). We observed that cells of *Mesotaenium* accumulated inclusions resembling LDs (Figure 6a) upon prolonged exposure to stress. Consistently, these globular structures were stained by BODIPY™ 493/503 (EM/EX), a common dye for lipid and oil-rich compartments (Listenberger and Brown, 2007; Kretzschmar et al., 2020). Under different conditions of temperature and light conditions, counts of LDs per cell showed significant differences (Figure 6b, Suppl. Table 8). We observed that the CGI-58 homolog was the 10th most connected hub in cluster green (Figure 5b). CGI-58 is key to lipid homeostasis, causing, if perturbed, the Chanarin-Dorfman syndrome in humans and LD overaccumulation in *Arabidopsis* (Lass et al., 2006; James et al., 2010; Figure 5c). Further, differential gene expression profiles pinpointed elevation of transcripts for characteristic LD protein homologs such as HSD1 and oleosin (OLE7) under high temperature and moderate light conditions (29 °C, 21 – 130 μmol photons m-2 s-1) and LD-associated protein (LDAP) and PUX10 under high temperature and light conditions (21-29 °C, 130 – 528 μmol photons m^−2^ s^−1^) (Figure 6c).

**Figure 6:**
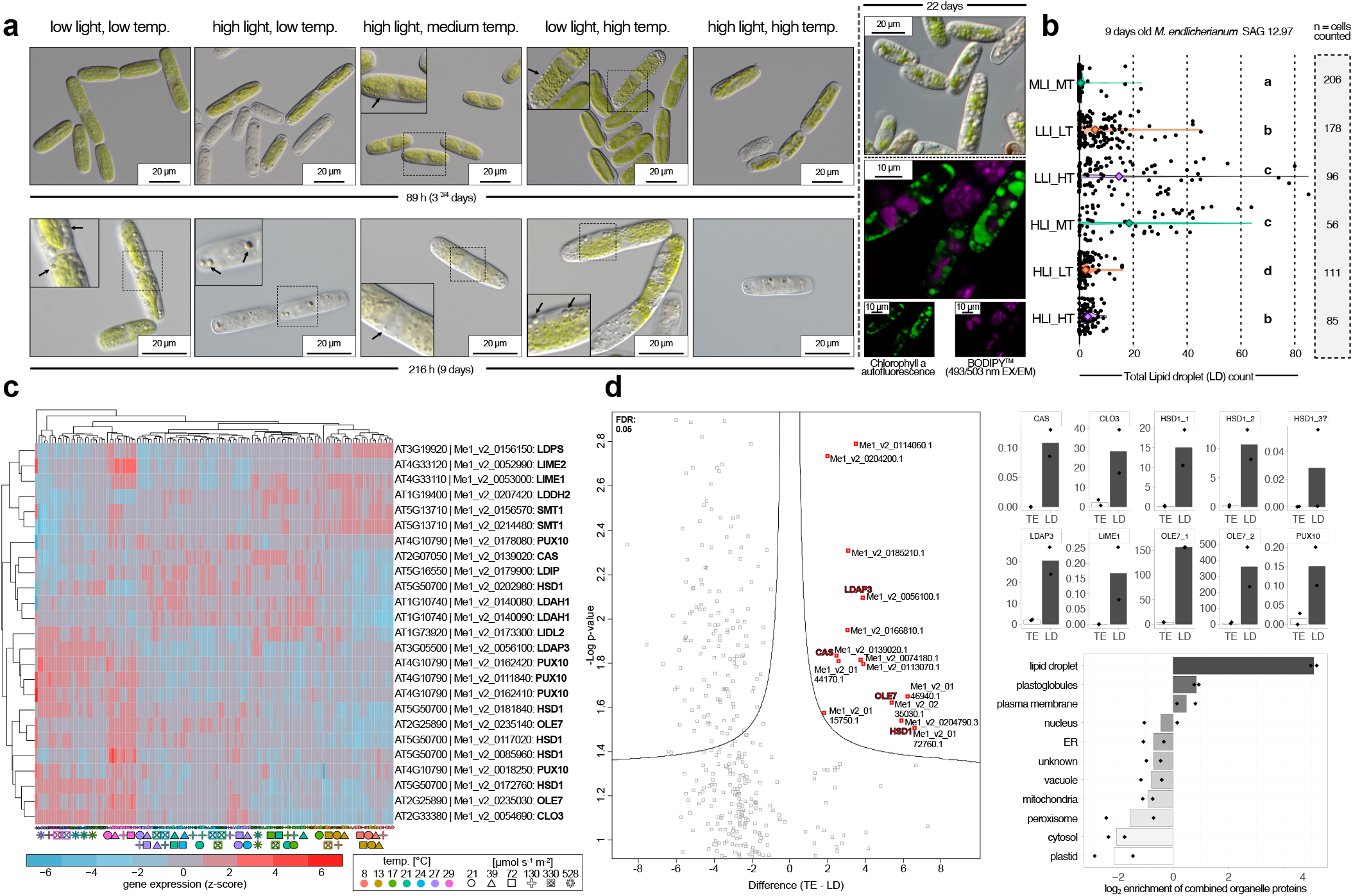
Lipid droplets accumulate in *Mesotaenium* upon changing environments. (a) Differential interference contrast (DIC) and confocal laser scanning micrographs of *Mesotaenium endlicherianum* SAG 12.97 cells accumulating lipid droplets (LDs; arrows). Cells were either subjected to different temperature/light conditions (see abbreviations below) of the gradient table for 89 h or 216 h. For confocal microscopy, algae were cultured independent of table conditions at 75 μmol photons m^−2^ s^−1^ and 22°C for 22 days. LDs are visible as distinct globular structures in NIC and were stained by the lipid stain BODIPY™ (false-colored green; 493 nm excitation; 503 nm emission) and chlorophyll autofluorescence (false-colored purple). (b) Violin plots of LD quantification after 9 days of exposure to different environmental conditions including statistical analysis using Mann-Whitney *U* statistics (significance grouping based on *p* value < 0.05; see also Suppl. Fig. 27) (c) Heat map of row-scaled z-scores of the expression of homologs for LD biogenesis and function (see also Suppl. Fig. 28). Temperature/light conditions are displayed at the bottom as symbols in different colors, *Arabidopsis thaliana* (AT) identifiers based on BLASTp search are shown on the right (d) Proteomic investigation into a lipid enriched phase extracted from *Mesotaenium endlicherianum* SAG 12.97 cells showing enrichment in hallmark proteins of LDs. Volcano plot showing significantly (FDR<0.05) enriched *Mesotaenium* proteins in the lipid enriched phase (right side) compared to proteins of the total extract (left side). Hallmark *A. thaliana* LD protein identifiers are annotated based on BLAST. Top bar on the left plot shows the relative, normalized iBAQ values for ten LD signature protein detected in *Mesotaenium*. Bottom bar plot shows the log2 enrichment of proteins characteristic for sub-cellular compartments. (LLI = Low light, LT = low temperature, MLI = moderate light, MT = moderate temperature, HLI = high light, HT = high temperature).

To scrutinize whether these structures are comparable to LDs of land plants, we performed sub-cellular fractionizations, obtained lipid-rich phases, and subjected them to proteomics using liquid chromatography-mass spectrometry (LC-MS). We identified 739 proteins in the putative LD fraction and 1574 proteins in the total extract (Suppl. Table 9). Of these, 14 were significantly enriched in the putative LD fraction (Figure 6 d, volcano plot) including hallmark LD proteins such as OLE, caleosin (CLO), steroleosin (HSD), and LDAP (Figure 6 d, bar plots). Overall, *Mesotaenium* responds to stress conditions by formation of LDs containing signature proteins for embryophytic LDs.

## DISCUSSION

Owing to their plain morphology, Zygnematophyceae emerged as unexpected closest algal relatives of land plants (Wickett et al., 2014; One Thousand Plant Transcriptomes Initiative, 2019; Hess et al., 2022). That said, the molecular programs of Zygnematophyceae speak of their close relationships to land plants. These point to a conserved chassis that likely operated in the last common ancestor of land plants and algae, featuring the proposed action of various hallmark genes (e.g., PYL homologs, GRAS family TFs and more) that were once considered land plant innovations. Building on the genomic resources for *Mesotaenium*, we have here delved into the molecular physiology and genetic programs of this alga, revealing which programs bear out when challenged with environmental cues.

Recent studies have proposed homology for the chassis of plastid–nucleus communication upon adverse environmental conditions between land plants and phragmoplastophytic streptophyte algae (Nishiyama et al., 2018; de Vries et al., 2018; Zhao et al., 2019). The GUN pathway likely has a conserved role in chloroplast transcription and streptophyte algal GUN1 homologs can rescue chloroplast retrograde signaling of Arabidopsis *Atgun1* mutants (Honkanen and Small, 2022); the degree of evolutionary conservation in the retrograde signaling pathway across streptophytes remains obscure (Honkanen and Small, 2022). Signals from damaged chloroplasts inhibit GLK1 expression in Arabidopsis (Martin et al., 2016). The negative correlation of module Blue (featuring *Me*GLK) with high light (leading to damaged chloroplasts) supports a role of *Me*GLK in operational retrograde signaling. Our data underscore that the wires between these components in plastid–nucleus communication are likely shared across more than 600 million years of streptophyte evolution and correlate with dealing with light regimes and adjustment of photosynthetic performance in the chloroplast also in the closest relatives of land plants.

One of the special features of plant and algal cells is their cell wall, forming their main interface with the environment. It is therefore not surprising that the cell wall is woven into a signaling network for environmental cues. In land plants, brassinosteroid-mediated signaling is part of a feedback loop for cell wall homeostasis and integrity (Wolf et al., 2014). While the involvement of brassinosteroids in streptophyte algae is doubtful—as is the case for many other phytohormones—our data suggest that there is a homologous chassis for a feedback loop for cell wall damage-based signaling that predates plant terrestrialization.

In land plants, the formation of LDs is known to occur under a variety of adverse environmental conditions (Gasulla et al., 2013; Mueller et al., 2015; Gidda et al., 2016). Stress-dependent formation of LDs likely evolved before land plants came to be (Li-Beisson et al., 2019; de Vries et al., 2020; de Vries and Ischebeck, 2020), but their molecular underpinnings outside of land plants remain unclear. Here, we confirmed the identity of these *Mesotaenium* LDs using confocal microscopy, LD-specific staining and proteomics. Our comprehensive transcriptomic data illuminate co-expressed modules that might constitute a homologous program for stress-dependent LDs that acted before plants conquered land.

## METHODS

### Algal culturing and gradient table setup

We used the axenic and genome-sequenced *Mesotaenium endlicherianum* SAG 12.97 (https://sagdb.uni-goettingen.de/detailedList.php?str_number=12.97) from the Algal Culture Collection, Göttingen, Germany (SAG, Friedl and Lorenz 2012, SAG). *Mesotaenium* was cultivated in C-Medium (Ichimura, 1971) for an average of 12 days in an aerated culture glass flasks (SCHOTT, Germany) at 80 μmol photons m^−2^ s^−1^. Prior to the experiment, cell density was analyzed using a LUNA™ Automated Cell Counter (Logos Biosystems, Annandale, VA, USA) and set to 2.03*10^7^ cells/ml (diluting with C-Medium if needed; settings for cell counting: Cell roundness: 60%, minimum size: 3 μm, maximum size: 60 μm), corresponding to Abs680nm = 0.33 (Epoch Microplatereader, BioTek Instruments, USA). For the gradient table setup algal suspension was distributed across 504 wells (42 twelve-well plates [tissue culture testplates 12 No. 92412, TPP, Switzerland]; 2.5 mL of culture per well). Plates were sealed with Surgical tape, Micropore™ tape (3M, Germany) to minimize evaporation. The 42 twelve-well plates were then placed on a table that generates a cross-gradient of temperature (8.6±0.5 °C to 29.2±0.5 °C on the x-axis) and irradiance (21.0±2.0 to 527.9±14.0 μmol photons m^−2^ s^−1^ on the y-axis) (see Suppl. Table 1). The temperature gradient was generated using a custom-made table (Labio, Czech Republic) equipped with true-daylight LEDs (sTube 2W 120 ver 11:11, Snaggi, Czech Republic) set to a 16:8 L/D cycle (Light from 6 am to 22 pm, Central European wintertime). *Mesotaenium* samples exposed to the 504 different conditions 65 hours (for sampling for RNAseq and physiological measurements) and 89 hours (for detailed light microscopy) on the gradient table. Condensed water at the top of the 12-well plates lids was removed three times in the 65 hours timespan by lightly tapping the lids twice.

### Plate reader

*In vivo* Abs480nm, Abs680nm, and Abs750nm of all 42 plates was measured after 65h exposition (4-6 hours after light on) with an absorbance microplate reader Epoch (BioTek Instruments, VT, USA). 9 data points per well were analyzed and averaged using software Gen5 2.0 (Biotek, USA), resulting in 108 measurements per 12-well plate per wavelength. For downstream analyses these values were averaged resulting in one value per 12-well plate per wavelength (Suppl. Fig. 1). After 89 h exposition 16 plates were chosen from the prominent gradients (the four most extreme conditions in the corners and a cross of vibrant growth along the two gradients) for analyzing a full absorption spectrum (300-900nm) using the same setup. (Suppl. Fig. 9, Suppl. Table 10).

### Photophysiological measurements

For maximum-quantum yield measurements (*F*_*v*_*/F*_*m*_) the maxi version of the IMAGING PAM (ImagMAX/L, M-series, Walz, Germany) with an IMAG-K5 CCD camera, controlled with the ImagingWinGigE (V2.32) software, was used. The *Mesotaenium* cultures in the 12-well plates were dark adapted for 10-30 min before measurement. Before measurements, the lid was removed. For the *F*_*v*_*/F*_*m*_ measurement a short saturation pulse (Intensity 3) was applied. The measurement settings on the IMAGING PAM were the following: measuring light 1, gain 3, damping 2, mean over AOI (area of interest) was turned off. No special SP-routine was applied to modify the signal to noise ratio of the chlorophyll fluorescence measurement.

### Statistical analysis of absorption and *F*_*v*_*/F*_*m*_ values and temperature/light cluster analysis

Statistical analysis of the absorption and the Fv/Fm values was done using Kruskal-Wallis test with post hoc test Fisher’s least significant difference (Conover, 1999) using R (version 4.1.3). P-values were Bonferroni corrected and grouped into significant groups using R packages ‘agricolae’ version 1.3-5 and ‘dplyr’ version 1.0.9. For heatmap generation of physiological values plotted against temperature/light R package ‘pheatmap’ version 1.0.12 was used. For cluster analysis the R package ‘factoextra’ version 1.0.7 was used. Clusters were generated using the eclust function with clustering function ‘kmeans’, with number of clusters set to six and for hierarchical clustering ‘euclidean’ was used as distance measure. Clusters were visualized with principal component analysis (PCA) in R.

### RNA extraction and sequencing

After absorption measurements, the twelve-well plates were put back on the table to let cells adjust to the table conditions again for a minimum of 5 minutes before harvesting them. For RNA extraction 0.4 mL were taken from every well of the 42 twelve-well plates on the table after pipetting the cells up and down twice to homogenize them. In total 4.8 mL liquid culture was taken per condition on the table (i.e., pooling 0.4 mL of each 12 wells per each of the 42 conditions). Samples were then centrifuged for 5 min at 20 °C and 4000 rpm. The supernatant was removed and the pellet was frozen at −80 °C. To extract RNA the Spectrum™ Plant Total RNA Kit (Sigma-Aldrich Chemie GmbH, Germany) was used according to the manufacturer’s instructions. For cell disruption samples in lysis buffer were ultrasonicated for 1 min and vortexed. RNA samples were treated with DNAse I (Thermo Fisher, Waltham, MA, USA) and shipped on dry ice to Novogene (Cambridge, UK) where they were quality checked with a Bioanalyzer (Agilent Technologies Inc., Santa Clara, CA, USA). Libraries were built based on total RNA using poly-T oligo-attached magnetic beads. Following fragmentation, synthesis of the first strand cDNA was carried out using random hexamer primers and second strand cDNA using dUTP, instead of dTTP. A directional size-selected library was built that included PCR-based amplification. Sequencing adapters were 5’ Adapter: 5’-AGATCGGAAGAGCGTCGTGTAGGGAAAGAGTGTAGATCTCGGTGGTCGCCGTATCATT-3’ and 3’ Adapter: 5’-GATCGGAAGAGCACACGTCTGAACTCCAGTCACGGATGACTATCTCGTATGCCGTCTTCT GCTTG-3’. The library was sequenced on an Illumina NovaSeq6000 platform.

### Quality control of reads

We checked the quality of our raw reads via FastQC (Andrews, Simon 2010) (v0.11.9) and summarized the results via MultiQC (Ewels et al. 2016) (v1.11). Based on these and the used adapter sequence, we filtered and trimmed reads via Trimmomatic (Bolger, Lohse, and Usadel 2014) (v 0.36) with these parameters: (“ILLUMINACLIP: novogene_adapter_sequences_Trimmomatic.fa:2:30:10:2:True LEADING:26 TRAILING:26 SLIDINGWINDOW:4:20 MINLEN:36”). We checked the quality of the trimmed reads with FastQC and MultiQC again.

### Genome annotation

The original annotation of *M. endlicherianum* (Cheng et al., 2019) had a lower number of genes compared to other Zygnematophyceae algae. We took advantage of our newly generated RNAseq dataset to improve genome annotation. Trimmed reads were mapped via HISAT2 (Pertea et al. 2016, 2) and assembled via StringTie (Pertea et al. 2016, 2). StringTie results showed many novel isoforms as well as novel transcripts. We also used BUSCO V5 (Manni et al. 2021) to measure the completeness of the gene models in annotation V1 independent of StringTie. Although the gene prediction method which used by BUSCO at the genome level is very efficient, it is not unexpected if it misses some proteins that were annotated in a genome via experimental, based on bioinformatic methods and NGS data, or ab-initio based gene prediction methods. Therefore, we expect that the BUSCO score based on the proteins of a gene model should be equal to or greater than the BUSCO score of the genome. When we compared the BUSCO score between the genome and protein sequences for *M. endlicherianum* with “viridiplantae.odb.10-2020-09-10”, we noticed that they show similar numbers (Suppl. Fig. 2). Therefore, we decided to re-annotate the genome of *M. endlicherianum* with our comprehensive RNA-Seq datasets as well as public protein and genome sequences published for its close relatives.

We annotated the *M. endlicherianum* genome using REAT (v0.6.1). Various gene models were predicted based on different types of evidence and methods. The final gene models and annotation V2 were based on agreement with the experimental evidence. At the end, we tried to quantify “completeness” and quality of the new annotation V2 and the old V1.

First, we used RNAseq evidence with REAT’s “Transcriptome Workflow” with HISAT2 (v2.2.1), Scallop (Shao and Kingsford 2017) (v0.10.5) and StringTie (v2.1.5). We also used Portcullis (Mapleson et al. 2018) (v1.2.4) to identify genuine junctions based on short reads alignments. This workflow uses Mikado (Venturini et al. 2018) (v2.3.4) to identify the “best” set of transcripts from multiple transcript assemblies.

Then, we used gene homology information from representative streptophytes in REAT’s “Homology Workflow”. SPALN (Gotoh 2008a; 2008b) (v2.4.7) was used to align representative protein sequences onto the *M. endlicherianum* genome. The representative dataset consisted on genome, gene models, and protein sequences of *Anthoceros agrestis* (Oxford strain) (Li et al. 2020), *Arabidopsis thaliana* (C.-Y. Cheng et al. 2017), *Azolla filiculoides* (Li et al. 2018), *Chara braunii* (Nishiyama et al. 2018), *Chlorokybus melkonianii* (Wang et al. 2020), *Chlamydomonas reinhardtii* (Merchant et al. 2007) (v5.6), *Klebsormidium nitens* (Hori et al. 2014), *Mesostigma viride* (Liang et al. 2019), *Marchantia polymorpha* (Montgomery et al. 2020) (v6.1r1), *Penium margaritaceum* (Jiao et al. 2020), *Physcomitrium patens* (Lang et al. 2018) (v3.3), *Selaginella moellendorffii* (Banks et al. 2011), and *Spirogloea muscicola* (S. Cheng et al. 2019). We also used the junction file produced by Portcullis. Since there were no close relatives of *M. endlicherianum* on the SPALN species-specific parameter set, we used three different closest possibilities (Angiosp, Chlospec, and MossWorts) and built three models. These alignments are filtered using a predefined set of criteria (*cf*. code on GitHub) including exon length, intron length, internal stop codon, among others. The final gene models of V2 were prepared by Mikado.

Afterwards, we used REAT’s “Prediction Workflow” to predict gene models *ab initio* and based on RNAseq and homology evidence. This uses Augustus (Stanke et al. 2006; Stanke, Tzvetkova, and Morgenstern 2006; Hoff and Stanke 2019) (v 3.4.0), SNAP (Korf 2004) (version 2006-07-28), Glimmer (Kelley et al. 2012) (v0.3.2), and CodingQuarry (Testa et al. 2015) (v2.0), which generate different gene models as the raw material for EvidenceModeler (Haas et al. 2008) (v1.1.1) that chooses the best set of exons and combine them in a gene model using weights (see GitHub) that could be adjusted for each sort of prediction and evidence. To include UTRs where possible, the EVM output is then processed by Mikado using UTR-containing gene models from the transcriptome and homology workflows as inputs, as well gene models classified by REAT as gold, silver, and bronze based on their agreement with the set of protein sequences from other streptophyte genomes (streptophyte algae and land plants), transcriptome alignment, homology alignment, and junctions. To train *ab initio* predictors, a user-defined number of models are randomly chosen in a user-defined ratio between (10%) mono-exonic and (90%) multi-exonic. These models were chosen from best classified models (gold and silver). For Augustus, we performed meta parameter optimization and train a model with kfold=8. Beside *ab initio* predictions, we used Augustus to predict gene models with three different weights for each evidence type as suggested by REAT authors (*cf*. code on GitHub).

At last, we used Minos (“Minos - a Gene Model Consolidation Pipeline for Genome Annotation Projects” [2019] 2022) which is gene model consolidation pipeline and produces external metrics based on DIAMOND “BLASTp/BLASTx” (Buchfink, Xie, and Huson 2015), Kallisto (Bray et al. 2016) (v0.46.2) expression quantification, coding potential calculator (CPC2 v0.1) (Kang et al. 2017, 2) and BUSCO assessments. These metrices pass through Mikado in combination with various gene models produced with different methods (as mentioned above), Minos determines the best gene model for each region based on user defined criteria (for details, see GitHub) and external metrics. Minos also put a tag on each gene model to categorize them based on a user defined threshold (we used default values) for sequence similarity coverage of homologs, BUSCO score, CPC score, TPM expression, and transcript score into “high confidence”, “low confidence”, and “predicted genes”.

### Genome annotation assessment

We used two methods to compare the quality of the new gene model with the published one. We compared the BUSCO scores of the annotated protein sequences as well as genome sequence using the reference “viridiplantae.odb.10-2020-09-10” dataset. We also used maker (Campbell et al. 2014) (v3.01.04) to calculate the AED (Eilbeck et al. 2009) to evaluate the agreement of the gene models with external evidences. Maker-P was used to build the *M. endlicherianum* gene model V1.

Further, we used the maker package to perform functional annotation via InterProScan and BLAST using agat (Dainat 2020) package (v0.9.2). Additionally, we performed a BLAST search against A. thaliana protein sequences (Araport11) and reported the best hit for each sequence in (Suppl. Table 11) and used eggNOGmapper (Huerta-Cepas et al. 2017; 2019) (v2.1.8) to perform functional annotation. We used DIAMOND (Buchfink, Xie, and Huson 2015) (v2.0.15) with ultra-sensitive mode, *e* value cutoff of 1e^−7^ and in an iterative manner. We used the protein sequences as our inputs and Viridiplantae (33090) as our taxonomy scope.

### RNA-Seq analysis: Pseudoalignment

In order to quantify gene expression, we used Kallisto (Bray et al. 2016) (v0.45.0). We indexed the transcriptome file with --kmer-size=31 parameter and used --bootstrap-samples 100 and --rf-stranded to quantify gene expression based on pseudoaligned reads. We used MultiQC to obtain an overview of alignment for each condition.

### Filtering, normalization, modeling mean-variance relationship, and data exploration

Kallisto quantification files were imported into R (v4.2.0) with tximport (Soneson, Love, and Robinson 2016) (v1.24.0) to calculate the counts from abundance via “lengthScaledTPM” based on our study design file (Suppl. Table 12). We used edgeR (Robinson, McCarthy, and Smyth 2010) (v3.38.1) for filtering and TMM-normalization (Robinson and Oshlack 2010) of the reads (genes with >1 count per million (CPM) at log2 scale in a least 3 samples—the number of replicates—were kept). Then, we used the voom function from limma (Ritchie et al. 2015; Phipson et al. 2016; Law et al. 2014; Liu et al. 2015) (v3.52.2) to model mean-variance relationship. The normalized expression table on the log2 scale is available in (Suppl. Table 13). We performed principal component analysis based on the expression table output of voom and visualized the result with ggplot2 (Ggplot2 n.d., 2) (v3.3.6). We visualized the heatmap of distance and Spearman correlation between all samples considering all genes via pheatmap (v1.0.12) and calculated clusters via the Euclidian method.

### RNA-Seq analysis: Weighted gene co-expression network analysis

We used WGCNA (Langfelder and Horvath 2008; 2012) package (v1.71) with the expression table produced by limma. We checked for and filtered out outliers as suggested by WGCNA authors (Suppl. Fig. 10). Then, we visualized the scale free topology model fit (R^2^) against the soft thresholds (ßs) to pick a ß for our network construction (Suppl. Fig. 11). We used signed network type and “bicor” as our correlation function for WGCNA. Based on these results, we picked 16 as our soft threshold ‘ß’. We experimentally chose a merging threshold if 0.25 after exploring different values from 0.2 to 0.4 and investigating the relationship between eigengenes and temperature, light intensity, *F*_*v*_*F*_*m*_, and absorption (Suppl. Fig. 12). We built the gene co-expression network using a merging threshold of 0.25 for modules, maximum portion of outliers as 0.05 and minimum module size of 30. Then, we visualized the correlation between each module eigengene and temperature, light intensity, *F*_*v*_*F*_*m*_, and absorption to identify which modules are more related to each treatment (Figure 4c). We provided a table for all genes, their module assignment, inter- and intramodular connectivity, gene significance for temperature and light intensity, correlation with temperature and light intensity, and their module membership (aka. Signed eigengene-based connectivity) in (Suppl. Table 5). We also visualized the graphical representation of the topological overlap matrix of our samples (Suppl. Fig. 13). In order to have a visual representation of gene expression in each module, we drew heatmaps for each module via pheatmap (using Euclidean method for calculating the distance and complete method clustering) (Suppl. Fig. 14). GO enrichment analysis was performed via clusterProfiler package (Yu et al. 2012; Wu et al. 2021) (v4.4.4) using the output of eggNOGmapper and adjusted p-value cut-off 0.05 and q-value cut-off of 0.05, considering only genes that are present in our GO term-to-gene table which was expressed and passed filtering as our background gene universe (Suppl. Table 6). Determining the proper background gene list has significant importance in enrichment analysis (Wijesooriya et al. 2022).

To see how *A. thaliana*’s well-known genes in stress-response mechanisms (downloaded from TAIR database via keyword search) were distributed across different modules we performed BLASTp searches against the new *M. endlicherianum* annotated proteins. We visualized the distribution of these IDs for different stress-related keywords in (Suppl. Fig. 15) and the expression of these genes across different samples via pheatmap (Suppl. Fig. 16). We defined as module hubs the top 20 genes (nodes) with the highest connectivity within each module (Suppl. Table 5 and 14).

### Differential gene expression analysis

We performed differential gene expression analysis using the limma package. We divided samples into multiple groups as follows: low light intensity (21 and 39 μmol photons m^−2^ s^−1^), medium light intensity (72 and 129 μmol photons m^−2^ s^−1^), high light intensity (329 and 527 μmol photons m^−2^ s^−1^), low temperature (8 °C and 12 °C), medium temperature (17 °C, 20 °C, and 23 °C), high temperature (26 °C and 29 °C; see grid/colored table layout in Figure 3). We performed all-against-all comparisons and an additional comparison of those samples from an *F*_*v*_*/F*_*m*_ < 0.5 versus low light intensity + medium temperature. We used duplicateCorrelation as suggested by Smyth et al. (2005) to consider technical replicates. We used clusterprofiler for GO-enrichment analysis (Wu et al. 2021) with adjusted p-value and q-value cutoff of 0.01 and only genes that were expressed and passed filtering as our background universe. The heatmap of gene expression profiles, dot plot and cnetplot of enriched GO-terms for each comparison is available in (Suppl. Table 14 and Suppl. Fig. 17 to 25).

### Phylogenetic analyses

We assembled a protein database based on the protein releases from the genomes of: *Anthoceros agrestis* BONN (Li et al., 2020), *Anthoceros punctatus* (Li et al., 2020), *Amborella trichopoda* (*Amborella* Genome Project, 2013), *Arabidopsis thaliana* (Lamesch et al., 2012), *Azolla filiculoides* (Li et al., 2018), *Bathycoccus prasinos* (Moreau et al., 2012), *Brassica oleracea* (Liu et al., 2014), *Brassica rapa* (Wang et al., 2010), *Brachypodium distachyon* (The International Brachypodium Initiative, 2010), *Capsella grandiflora* (Slotte et al., 2013), *Chara braunii* (Nishiyama et al., 2018), *Chlorokybus atmophyticus* (Wang et al., 2020), *Chlamydomonas reinhardtii* (Merchant et al., 2007), *Coccomyxa subellipsoidea* (Blanc et al., 2012), *Gnetum montanum* (Wan et al., 2018), *Klebsormidium nitens* (Hori et al., 2014), *Marchantia polymorpha* (Bowman et al., 2017), *Mesostigma viride* (Wang et al., 2020), *Micromonas pusilla, Micromonas* sp. (Worden et al., 2009), *Oryza sativa* (Ouyang et al., 2007), *Picea abies* (Nystedt et al., 2013), *Physcomitrium patens* (Lang et al., 2018), *Salvinia cucullata* (Li et al., 2018), *Selaginella moellendorffii* (Banks et al., 2011), *Solanum lycopersicum* (The Tomato Genome Consortium, 2012), *Theobroma cacao* (Argout et al., 2011), *Mesotaenium endlicherianum* (Cheng et al., 2019), *Ostreococcus lucimarinus* (Palenik et al., 2007), *Penium margaritaceum* (Jiao et al., 2020), *Spirogloea muscicola* (Cheng et al., 2019), *Ulva mutabilis* (De Clerck et al., 2018), *Volvox carteri* (Prochnik et al., 2010).

Homologs for proteins were detected using BLASTp with Arabidopsis and *Mesotaenium* proteins as query against the aforementioned proteins as database. Alignments were computed using MAFFT v7.490 (Katoh and Standley, 2013). All phylogenies were computed with IQ-TREE multicore version 1.5.5 (Nguyen et al., 2015); their respective best model for protein evolution was determined using ModelFinder (Kalyaanamoorthy et al., 2017) according to Bayesian Information Criterion and 1000 ultrafast bootstrap replicates; 1000 ultrafast bootstrap replicates (Hoang et al., 2018) were carried out and 100 Felsenstein bootstraps (Felsenstein, 1985) for the LDAP phylogeny.

### Differential interference contrast and confocal laser scanning microscopy

Differential interference contrast (DIC) imaging was done for all replicates from the table with a Olympus BX-60 microscope (Olympus, Japan) with a ProgRes C14plus camera and the ProgRes® CapturePro Software (version 2.9.01) (JENOPTIK AG, Jena, Germany). The morphology of chosen conditions (see Supplemental Figure 1) of *Mesotaenium* cells that were 89 h on the table was analyzed.

For algae that were used for quantifying the abundance of lipid droplet per cell, a ZEISS Axioscope 7 microscope (Carl Zeiss, Germany) was used including the _ZEN_ software (Carl Zeiss, Germany). Lipid droplet count was carried out in FIJI (Schindelin et al., 2012). For statistical analysis of the lipid droplet count data, we first used a Shapiro-Wilk test (Shapiro and Wilk, 1965) to assess normality and used Mann-Whitney U tests (Mann and Whitney, 1947) with R (version 3.6.1) accordingly.

Confocal laser scanning microscope was done on a Zeiss LSM780 (Carl Zeiss) set as in Müller et al. (2017). For the staining of the LD structures, we used the neutral lipid specific stain BODIPY™ 493/503 (EM/EX) (Merck). *Mesotaenium* cells were grown for 22 days on WHM-medium at 70-80 μmol photons m^−2^ s^−1^ and 22°C. These cells were ultrasonicated for 1 min with 1:500 BODIPY and incubated on a shaker for 5 min before visualization.

### Lipid droplet isolation and proteomics

For lipid droplet isolation 23 days old *Mesotaenium* cells grown on WHM-Medium at 70-80 μmol photons m^−2^ s^−1^ and 22 °C were homogenized using a Tenbroeck or potter homogenizer in lipid droplet isolation buffer (10 mM sodium phosphate buffer pH 7.5, 200 μM PMFS, 0.5 mM DSP, 10 mM N-Ethylmaleimide). The resulting centrifuged supernatant of a 100 x g spin for 1 min was considered as total extract (TE). After two further high speed centrifugations (SW40 Ti for 1h, 4°C at 100000 x *g*, TLA120 for 1h at 100000 x *g* and 4°C) the floating fat pad was precipitated at −20 °C using 100% ethanol overnight. The precipitated pellet was washed with 80% ethanol twice, dried and then suspended in 6M urea. Protein concentration was determined using BCA. An in-gel SDS gel digestion was done with trypsin adapted from Shevchenko et al. (1996). C18 Stage tip purification was done according (Rappsilber et al., 2003; 2007). Protein samples were analyses using LC-MS. For this, peptide samples were reconstituted in 20 μl LC-MS sample buffer (2% acetonitrile, 0.1% formic acid). 2 μl of each sample were subjected to reverse phase liquid chromatography for peptide separation using an RSLCnano Ultimate 3000 system (Thermo Fisher Scientific). Therefore, peptides were loaded on an Acclaim PepMap 100 pre-column (100 μm x 2 cm, C18, 5 μm, 100 Å; Thermo Fisher Scientific) with 0.07% trifluoroacetic acid at a flow rate of 20 μL/min for 3 min. Analytical separation of peptides was done on an Acclaim PepMap RSLC column (75 μm x 50 cm, C18, 2 μm, 100 Å; Thermo Fisher Scientific) at a flow rate of 300 nL/min. The solvent composition was gradually changed within 94 min from 96 % solvent A (0.1 % formic acid) and 4 % solvent B (80 % acetonitrile, 0.1 % formic acid) to 10 % solvent B within 2 minutes, to 30 % solvent B within the next 58 min, to 45% solvent B within the following 22 min, and to 90 % solvent B within the last 12 min of the gradient. All solvents and acids had Optima grade for LC-MS (Thermo Fisher Scientific). Eluting peptides were on-line ionized by nano-electrospray (nESI) using the Nanospray Flex Ion Source (Thermo Fisher Scientific) at 1.5 kV (liquid junction) and transferred into a Q Exactive HF mass spectrometer (Thermo Fisher Scientific). Full scans in a mass range of 300 to 1650 m/z were recorded at a resolution of 30,000 followed by data-dependent top 10 HCD fragmentation at a resolution of 15,000 (dynamic exclusion enabled). LC-MS method programming and data acquisition was performed with the XCalibur 4.0 software (Thermo Fisher Scientific). Afterwards the raw proteome data were analyzed using Max Quant software version 1.6.2.10 (Cox and Mann, 2008). The database for this analysis was our new V2 gene model data. The data were then further processed by the Perseus (1.6.2.2) software (Cox et al., 2008; Tyanova et al., 2016).

## Data availability

All RNAseq reads have been uploaded to NCBI SRA and can be accessed under Bioproject PRJNA832564 and SRA accessions SRR18936040 to SRR18936170. Codes and Data used for genome re-annotaiton, WGCNA and differential gene expression analysis are available on our GitHub page https://github.com/deVries-lab/Response_to_a_gradient_of_environmental_cues_in_mesotaenium_endlicherianum. Proteomic data have been uploaded to PRIDE. Furthermore, data can be interactively explored at https://mesotaenium.uni-goettingen.de

## ACKNOWLEDGEMENT

We thank René Heise for excellent technical support. J.d.V. thanks the European Research Council for funding under the European Union’s Horizon 2020 research and innovation programme (Grant Agreement No. 852725; ERC-StG “TerreStriAL”). J.d.V., U.H., and H.B. are grateful for support through the German Research Foundation (DFG) within the framework of the Priority Programme “MAdLand – Molecular Adaptation to Land: Plant Evolution to Change” (SPP 2237; VR 132/4-1; BU 2301/6-1), in which T.R. is a PhD student and A.D., J.M.R.F.-J, and I.I. partake as associate members. A.D. is grateful for being supported through the International Max Planck Research School (IMPRS) for Genome Science. J.M.R.F.-J. and T.R. gratefully acknowledge support by the Ph.D. program “Microbiology and Biochemistry” within the framework of the “Göttingen Graduate Center for Neurosciences, Biophysics, and Molecular Biosciences” (GGNB) at the University of Goettingen. P.S. was supported by the GGNB in frame of the PRoTECT program at the University of Goettingen. T.I. acknowledges funding from the Deutsche Forschungsgemeinschaft (DFG; GRK 2172-PRoTECT). M.M. is supported by Singaporean Ministry of Education grant T2EP30122-0001. P.S. is grateful for support from the Studienstiftung des Deutschen Volkes. We thank Prof. Dr. Christiane Gatz and Dr. Guido Kriete for giving us access to the ImagMAX/L PAM in the Department of Plant Molecular Biology and Physiology.

## CONTRIBUTIONS

J.d.V. and M.L. conceived the project. J.d.V. coordinated the project with M.M. M.L. provided plant materials. J.M.R.F.-J., T.D., and T.R. performed experimental work. A.D. carried out computational analysis. O.V., J.M.R.F.-J., P.S., T.I., D.K. and G.H.B. performed proteomics. H.B. investigated cell division patterns. M.H. and U.H. investigated photomorphogenesis patterns. A.D. and R.S. built web resources. J.d.V., A.D., and J.M.R.F.-J. contributed to writing the manuscript. J.d.V. organized the manuscript. All authors commented, discussed, and provided input on the final manuscript.

## SUPPLEMENTAL FIGURES

Supplemental Figure 1. *F*_*v*_*/F*_*m*_ and absorption values of all replicates of gradient tables; representative micrographs of the most extreme corners and of vividly growing algae along the two gradients.

Supplemental Figure 2. BUSCO comparison between genome, protein sequences V1, protein sequences V2

Supplemental Figure 3. Cumulative fraction of annotation vs AED plot for gene model V1 and V2

Supplemental Figure 4. Module membership versus Gene Significance for genes in different modules with respect to *F*_*v*_*/F*_*m*_

Supplemental Figure 5. Module membership versus Gene Significance for genes in different modules with respect to Temperature

Supplemental Figure 6. Module membership versus Gene Significance for genes in different modules with respect to light intensity

Supplemental Figure 7. Heatmap of the correlation between module eigengenes and light intensity, temperature, absorption, replicate, and *F*_*v*_*/F*_*m*_ as well as student test p-value

Supplemental Figure 8. The GLK alignment

Supplemental Figure 9. Absorption spectra of all replicates at chosen conditions.

Supplemental Figure 10. Sample dendrogram and trait heatmap to identify outliers for WGCNA

Supplemental Figure 11. Picking a soft threshold for WGCNA based on scale independence and Mean connectivity

Supplemental Figure 12. Clustering of different modules and traits based for identifying a merging threshold

Supplemental Figure 13. The graphical representation of the topological overlap matrix

Supplemental Figure 14. Heatmap of gene expression Z-score values for each module

Supplemental Figure 15. Distribution of best blast hit of *A. thaliana* stress response genes among WGCNA modules

Supplemental Figure 16. Heatmap of best blast hit of *A. thaliana* stress response genes in *M. endlicherianum* across different growth conditions

Supplemental Figure 17. Dotplot, cnetplot and heatmaps of DEGs comparing FvFm control vs stress

Supplemental Figure 18. Dotplot, cnetplot and heatmaps of DEGs comparing HLI_HT vs LLI_MT

Supplemental Figure 19. Dotplot, cnetplot and heatmaps of DEGs comparing MLI_HT vs LLI_MT

Supplemental Figure 20. Dotplot, cnetplot and heatmaps of DEGs comparing LLI_HT vs LLI_MT

Supplemental Figure 21. Dotplot, cnetplot and heatmaps of DEGs comparing HLI_MT vs LLI_MT

Supplemental Figure 22. Dotplot, cnetplot and heatmaps of DEGs comparing MLI_MT vs LLI_MT

Supplemental Figure 23. Dotplot, cnetplot and heatmaps of DEGs comparing HLI_LT vs LLI_MT

Supplemental Figure 24. Dotplot, cnetplot and heatmaps of DEGs comparing MLI_LT vs LLI_MT

Supplemental Figure 25. Dotplot, cnetplot and heatmaps of DEGs comparing MLI_LT vs LLI_MT

Supplemental Figure 26. Fully-labeled phylogenies of hub genes.

Supplemental Figure 27. Lipid droplet count setup 2.

Supplemental Figure 28. LDAP phylogeny.

## SUPPLEMENTAL TABLES

Supplemental Table 1. Temperature and light intensity measurements of all 504 coordinates on the gradient table.

Supplemental Table 2. All 504 *F*_*v*_*/F*_*m*_ and absorption measurements of all replicates.

Supplemental Table 3. Number of genes and transcripts in gene model V2

Supplemental Table 4. The general stats of raw reads, trimmed reads, and pseudoalignment

Supplemental Table 5. Summary of WGCNA Results

Supplemental Table 6. The results of GO-enrichment analysis for all modules of WGCNA

Supplemental Table 7. The list of top20 hubs for each module.

Supplemental Table 8. Counts of lipid droplets in micrographs.

Supplemental Table 9. Full proteomic results, showing Mesotaenium gene model V2 identifiers, Arabidopsis gene identifiers, and IBAQ values.

Supplemental Table 10. All data on absorption spectra of all replicates at chosen conditions.

Supplemental Table 11. Best blast hit of *M. endlicherianum* gene model against *A. thaliana* (Araport11)

Supplemental Table 12. Study design file used for RNASeq analysis

Supplemental Table 13. The CPM normalized expression table on the log2 scale

Supplemental Table 14. The GO-enrichment results of 9 pairwise comparisons

