## Supplemental Figures S1 to S28 for "Environmental gradients reveal stress hubs predating plant terrestrialization"

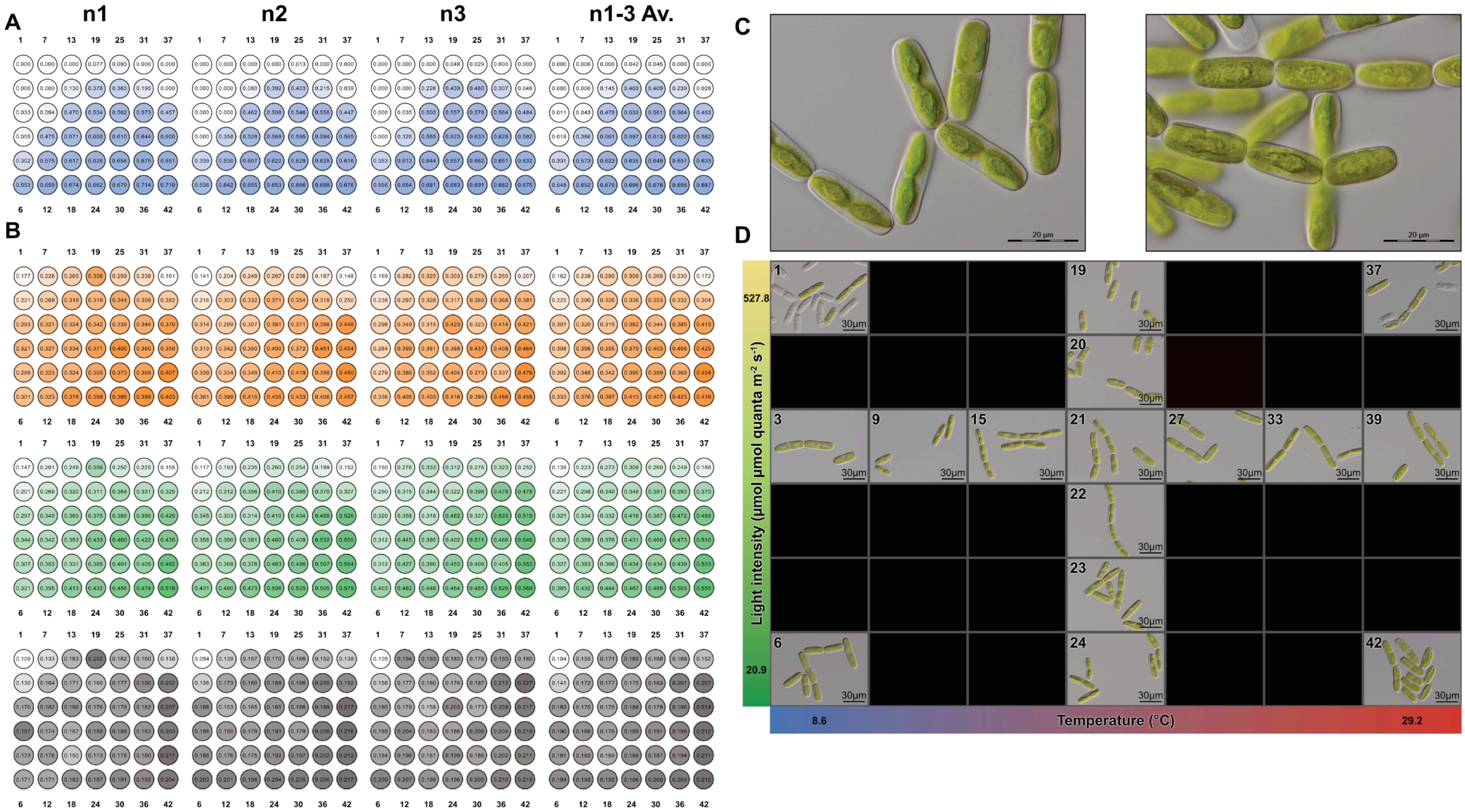


**Supplementary Figure S1: Fv/Fm, Absorption, Morphology. (a)** Fv/Fm values of all biological replicates (n1-n3) as well as the averaged values (n1-3 Av.). Blue color gradient corresponds to increase/decrease of values. Numbers shown at the top/bottom indicate 12-well plate numbering on the gradient table. **(b)** Absorption values of all biological replicates (n1-n3) as well as the averaged values (n1-3 Av.). Color gradient corresponds to increase/decrease of values. Colors indicate measured wavelength: orange gradients = Absorption measured at λ 480 nm, green gradients = Absorption measured at λ 680 nm, grey gradients = Absorption measured at λ 750 nm. Numbers shown at the top/bottom indicate 12-well plate numbering on the gradient table. **(c)** Differential interference contrast (DIC) micrographs of SAG 12.97 cells grown on C-Medium (growth conditions see methods: growth conditions prior to exposure to environmental conditions.) **(d)** Differential interference contrast (DIC) micrographs of SAG 12.97 under most extreme environmental conditions (four corners: samples 1,6,37,42) as well as along an irradiance gradient at 21°C (samples 19-24) and a temperature gradient at 130 µmol photons $m^{-2}$ $s^{-1}$ (samples 3,9,15,21,27,33,39).

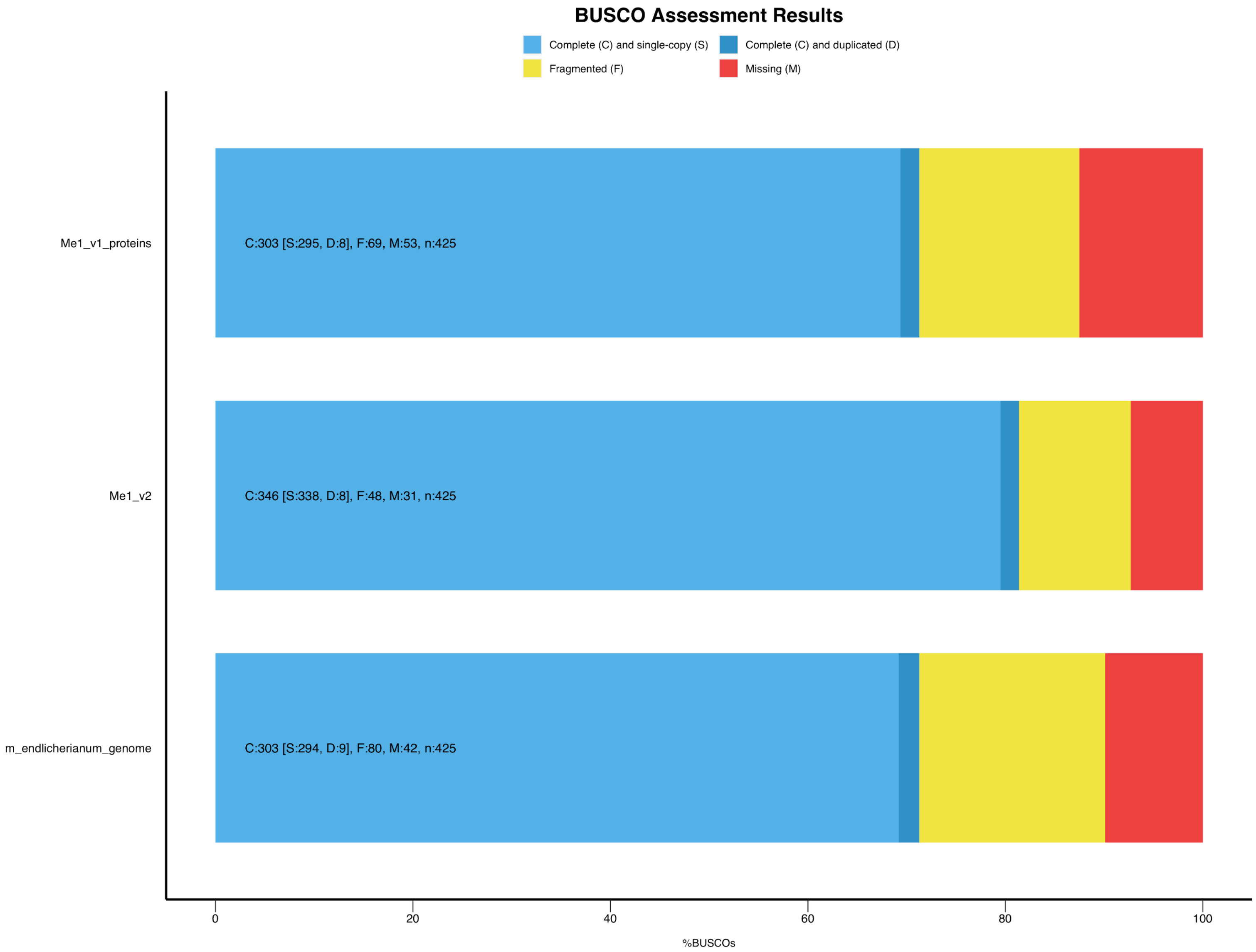


**Supplementary Figure S2: Comparison of BUSCO scores.** Comparisosns were carried out between genome, protein sequences V1, and protein sequences V2. The BUSCO score of annotation V2 resulted in less missing (red), less fragmented (yellow) and more complete (blue) genes.

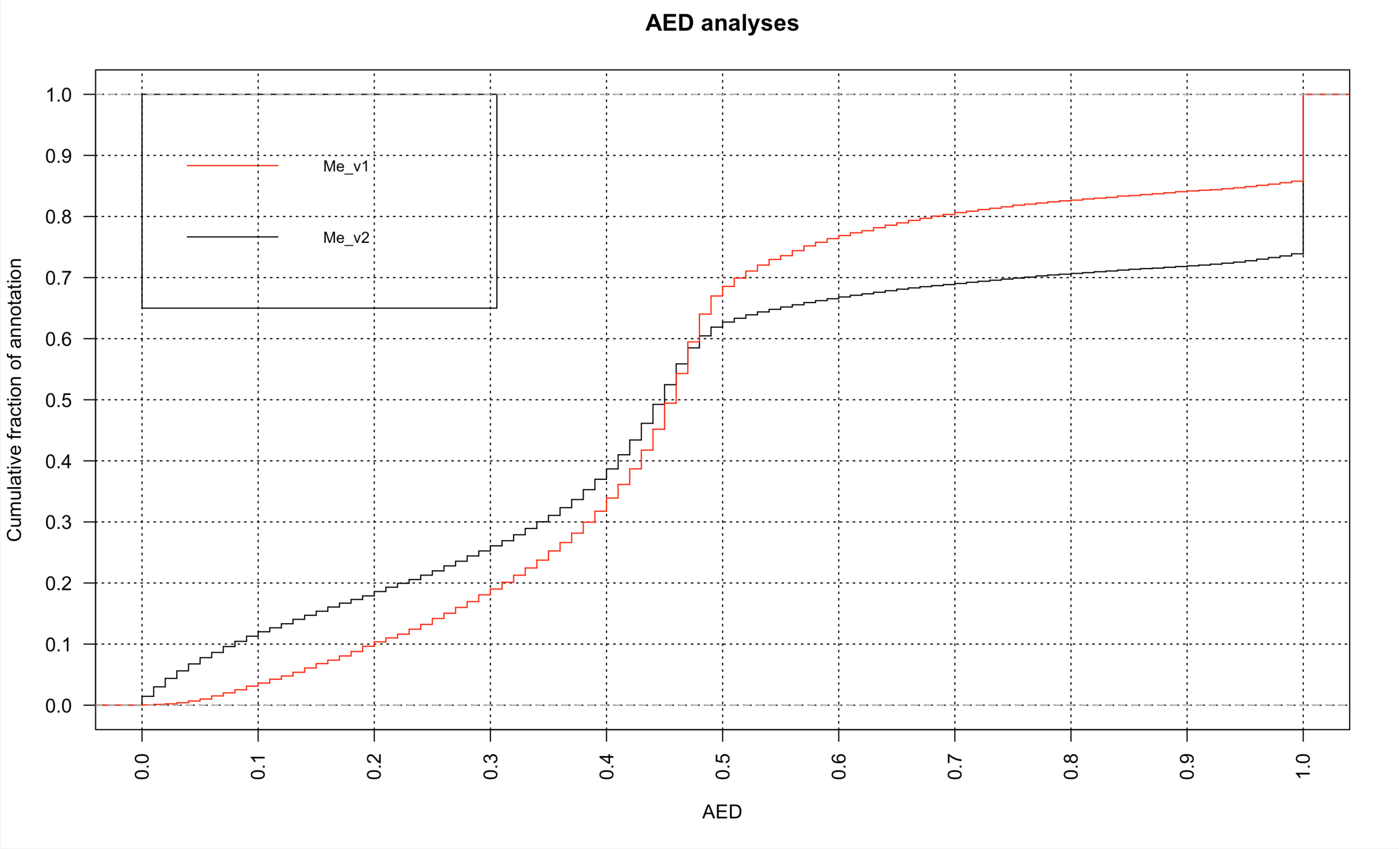


**Supplementary Figure S3: Quality assessment of the annotations.** Cumulative fraction of annotation versus annotation edit distance (AED) plot for annotations V1 and V2. The red line marks V1, the black line marks V2.

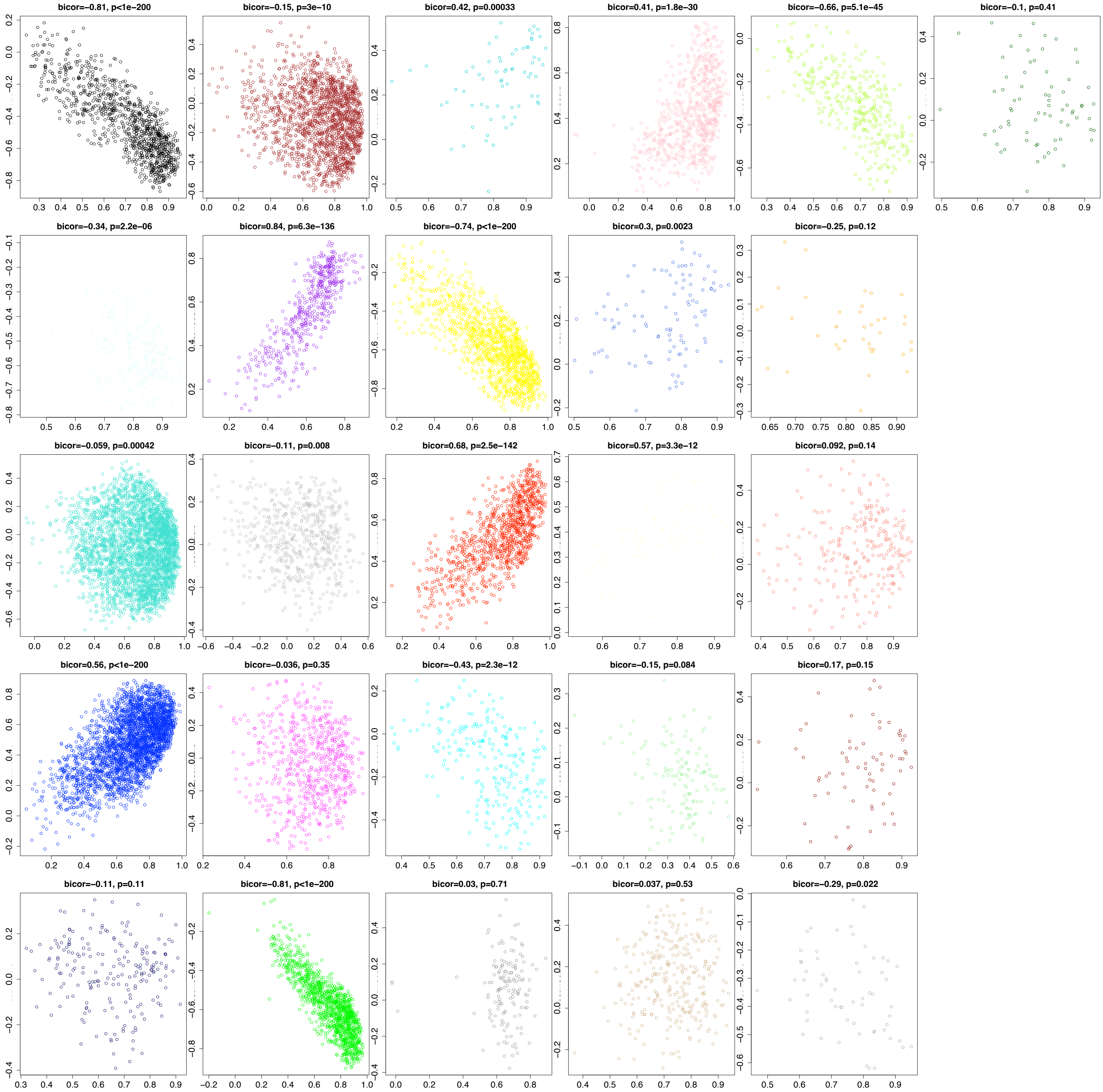


**Supplementary Figure S4: Module membership versus Gene Significance for genes in different modules with respect to Fv/Fm.** Colors correspond to the module name. X-axis shows "Module Membership in the module" and Y-axis shows "Gene significance for trait (Fv/Fm)".

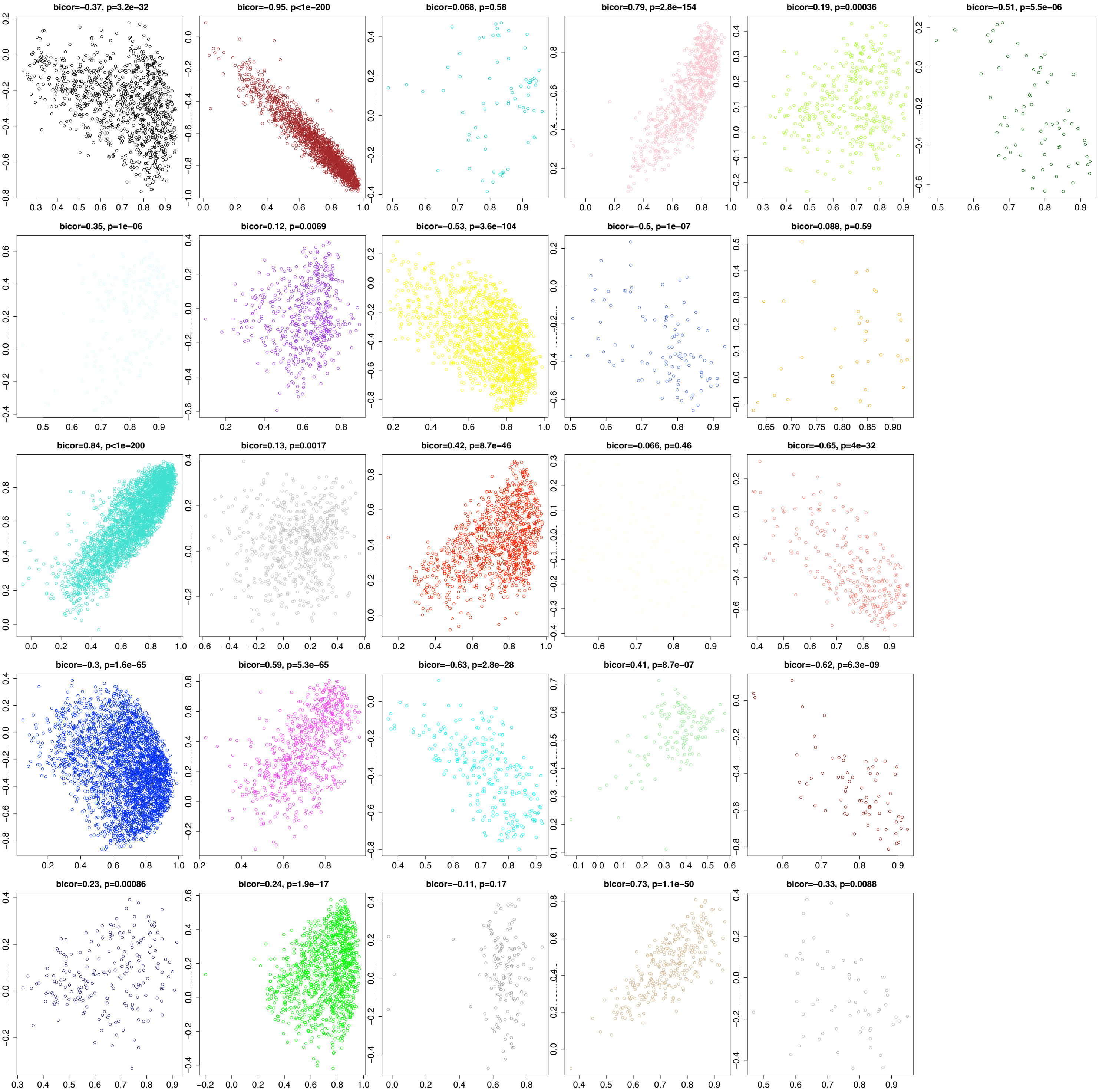


**Supplementary Figure S5: Module membership versus Gene Significance for genes in different modules with respect to Temperature.** Colors correspond to the module name. X-axis shows "Module Membership in the module" and Y-axis shows "Gene significance for trait (Temperature)".

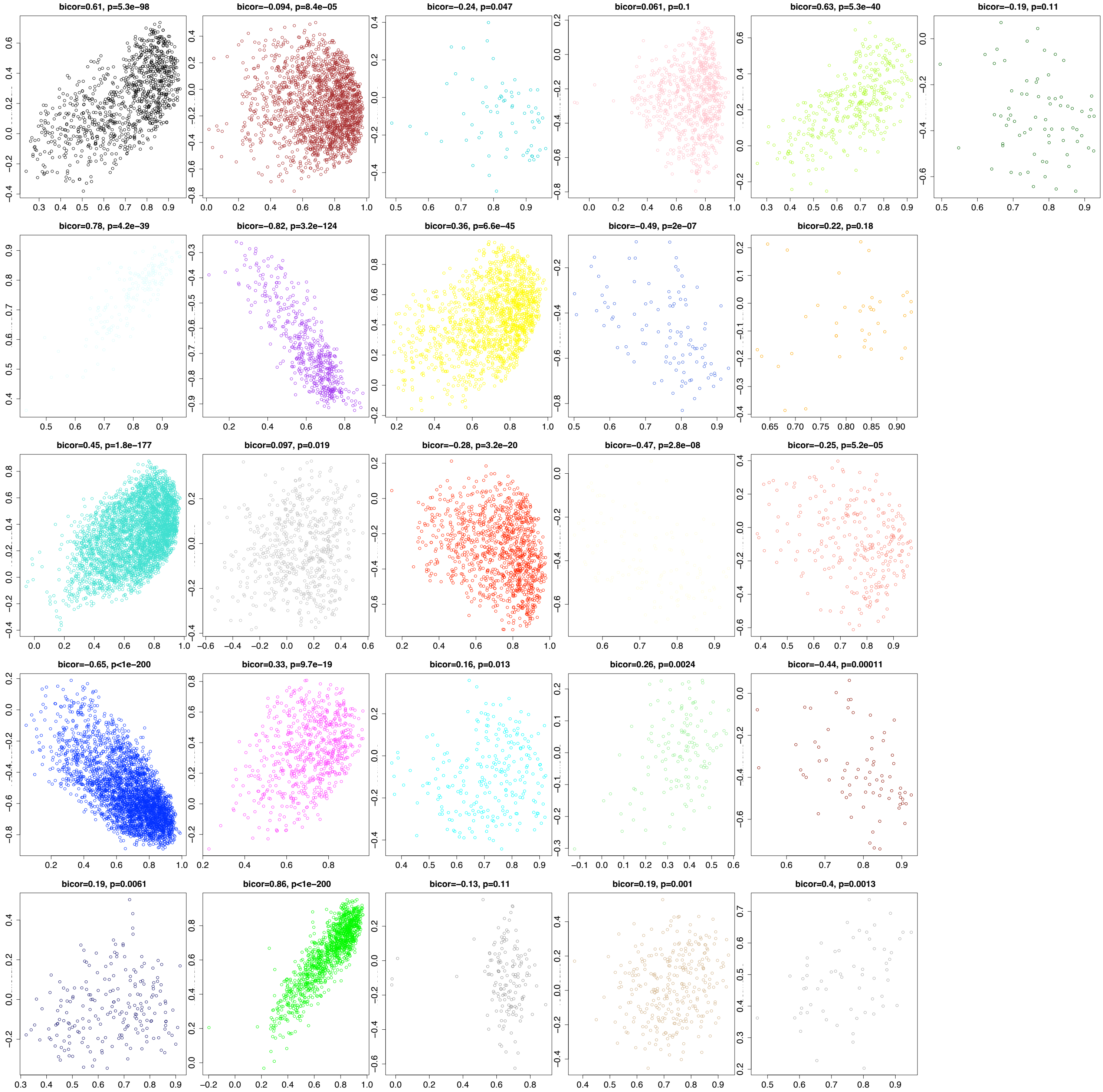


**Supplementary Figure S6: Module membership versus Gene Significance for genes in different modules with respect to light intensity.** Colors correspond to the module name. X-axis shows "Module Membership in the module" and Y-axis shows "Gene significance for trait (light intensity)".

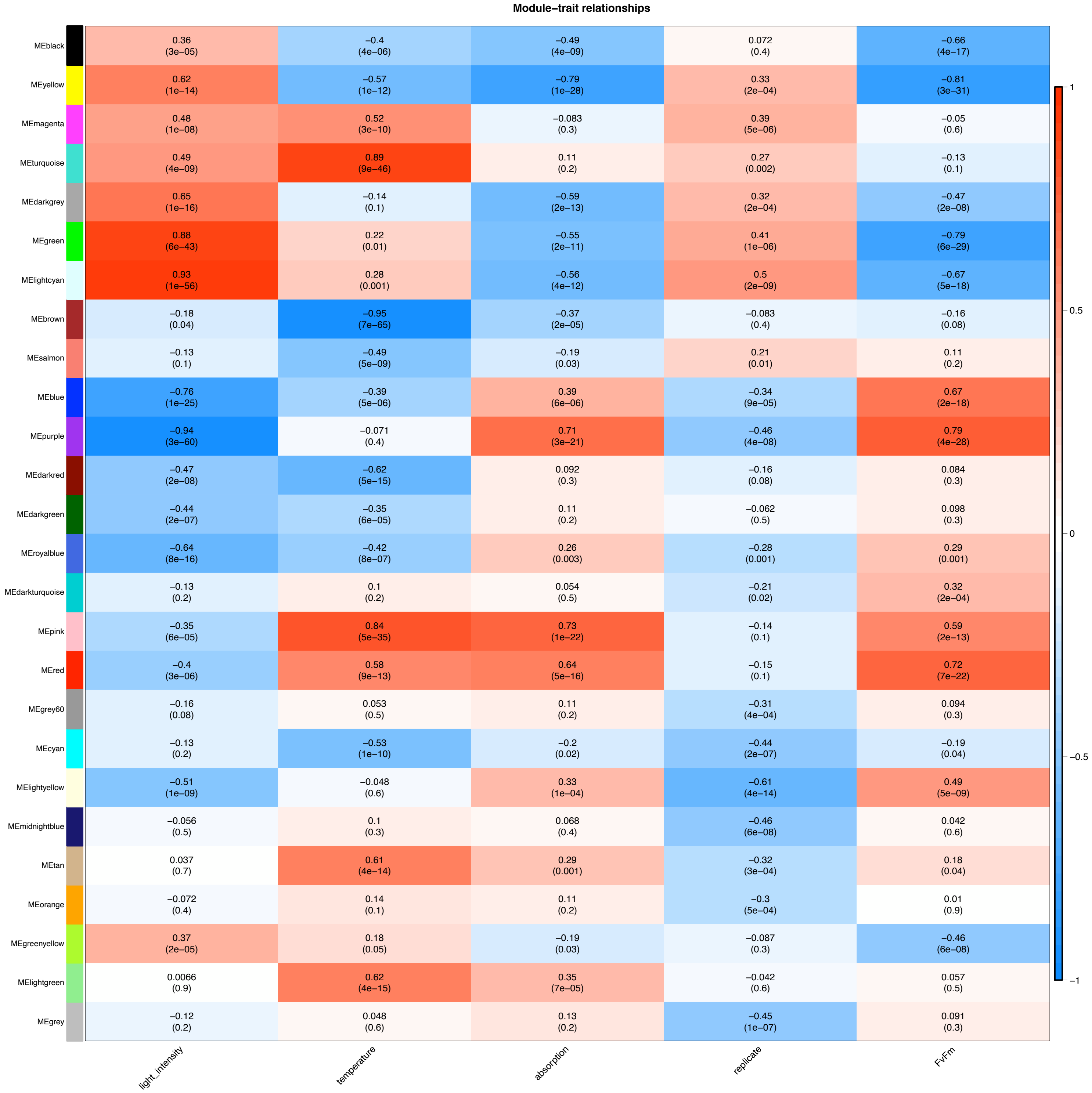


**Supplementary Figure S7: Module–trait correlation based on eigengenes.** Heatmap of the correlation between module eigengene expression behaviour and the parameters light intensity, temperature, absorption, replicate, and Fv/Fm as well as student test p-value. Corresponding to the main Figure 4b.

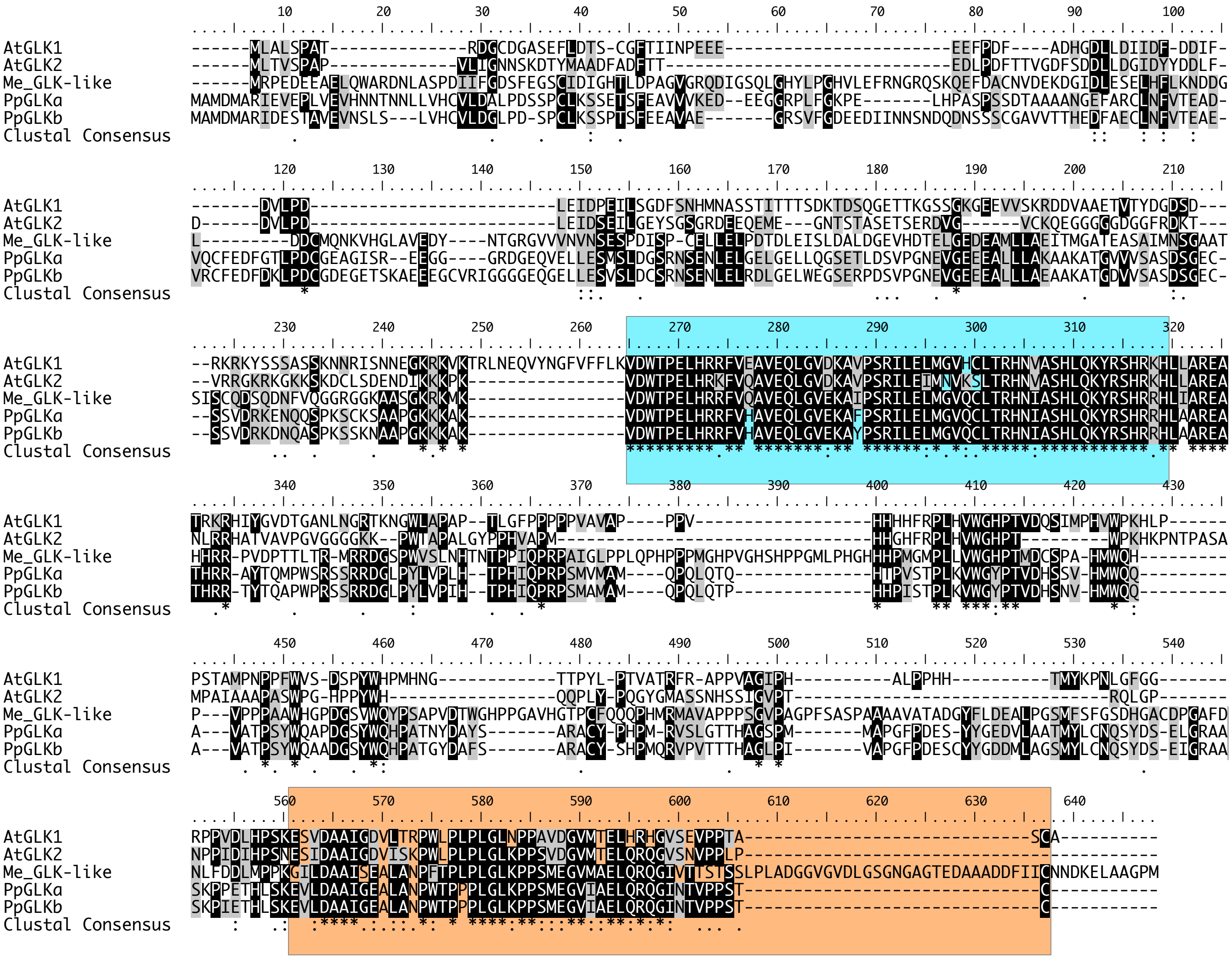

**Supplementary Figure S8: Alignment Golden2-like homologs and domains.** GLK homologs of Arabidopsis (At), *Mesotaenium endlicherianum* (Me), *Physcomitrium patens* (Pp) were aligned and a consensus was computed. Teal labels the DBD region, orange the GCT box.

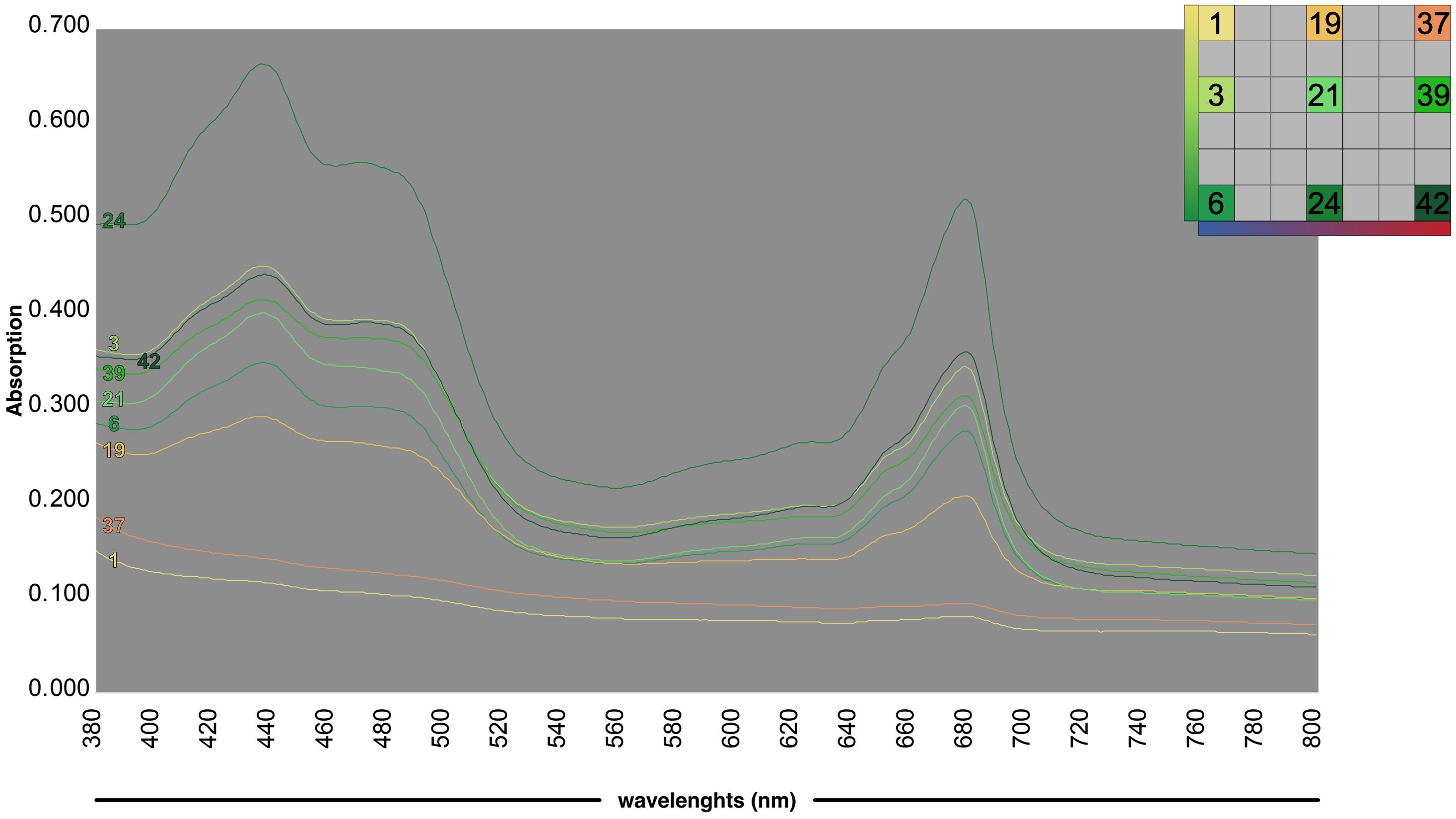


**Supplementary Figure S9: Full Absorption Spectra.** Shown are measured absorption spectra measured from λ 380 nm to λ 800 nm. Averaged values of three biological triplicates are shown (see individual values in Suppl. Table 10). Measured samples were taken from the most extreme environmental conditions (samples 1, 6, 37, 42) as well as along the whole temperature gradient at 130 µmol photons $m^{-2}$ $s^{-1}$ (samples 3, 21, 39) and along the whole irradiance gradient at 21°C (samples 24, 21, 19).

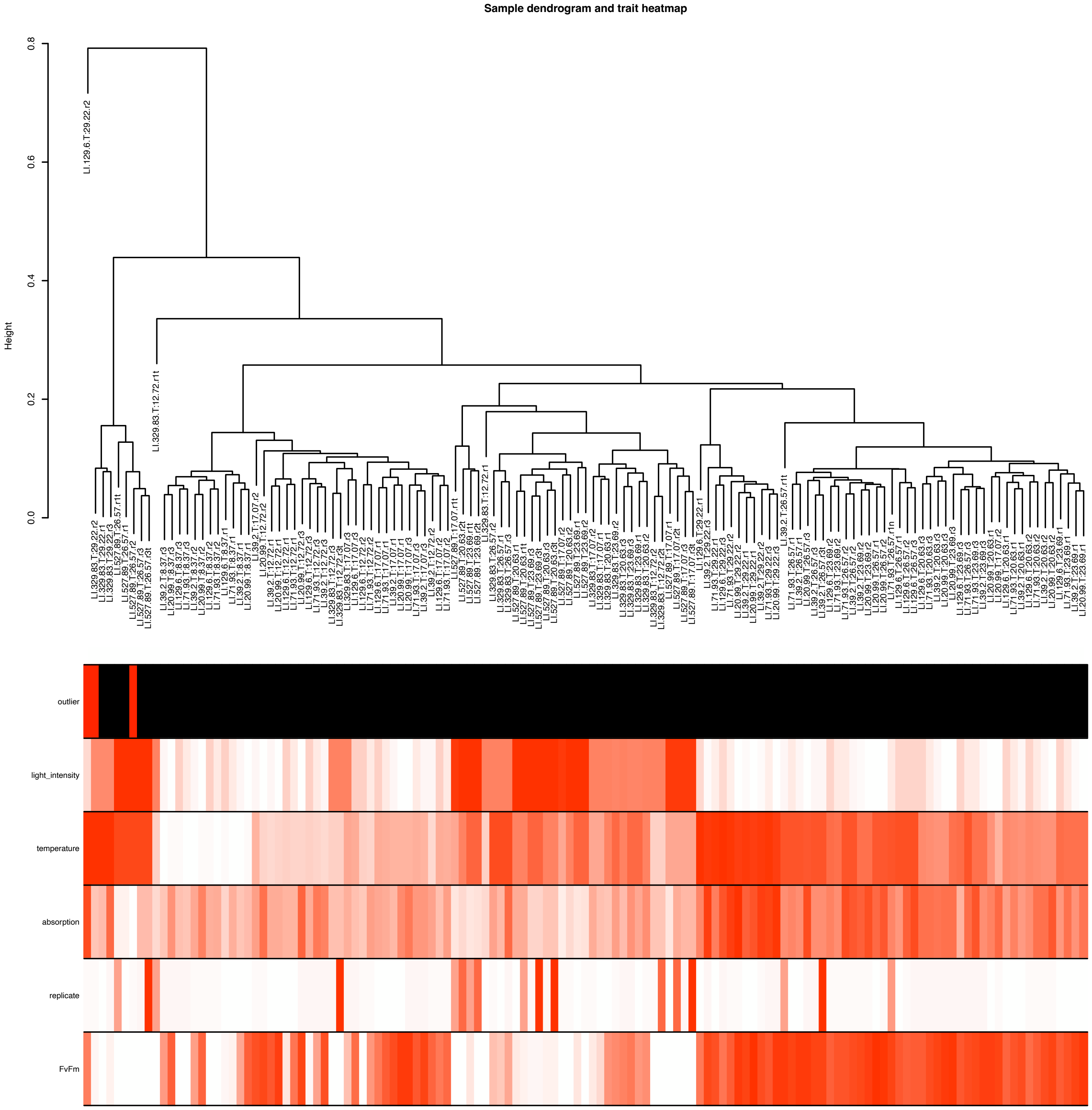


**Supplementary Figure S10: Samples analysed in WGCNA. A sample dendrogram was computed (top) and projected on top of a trait heatmap to identify outliers for WGCNA.**

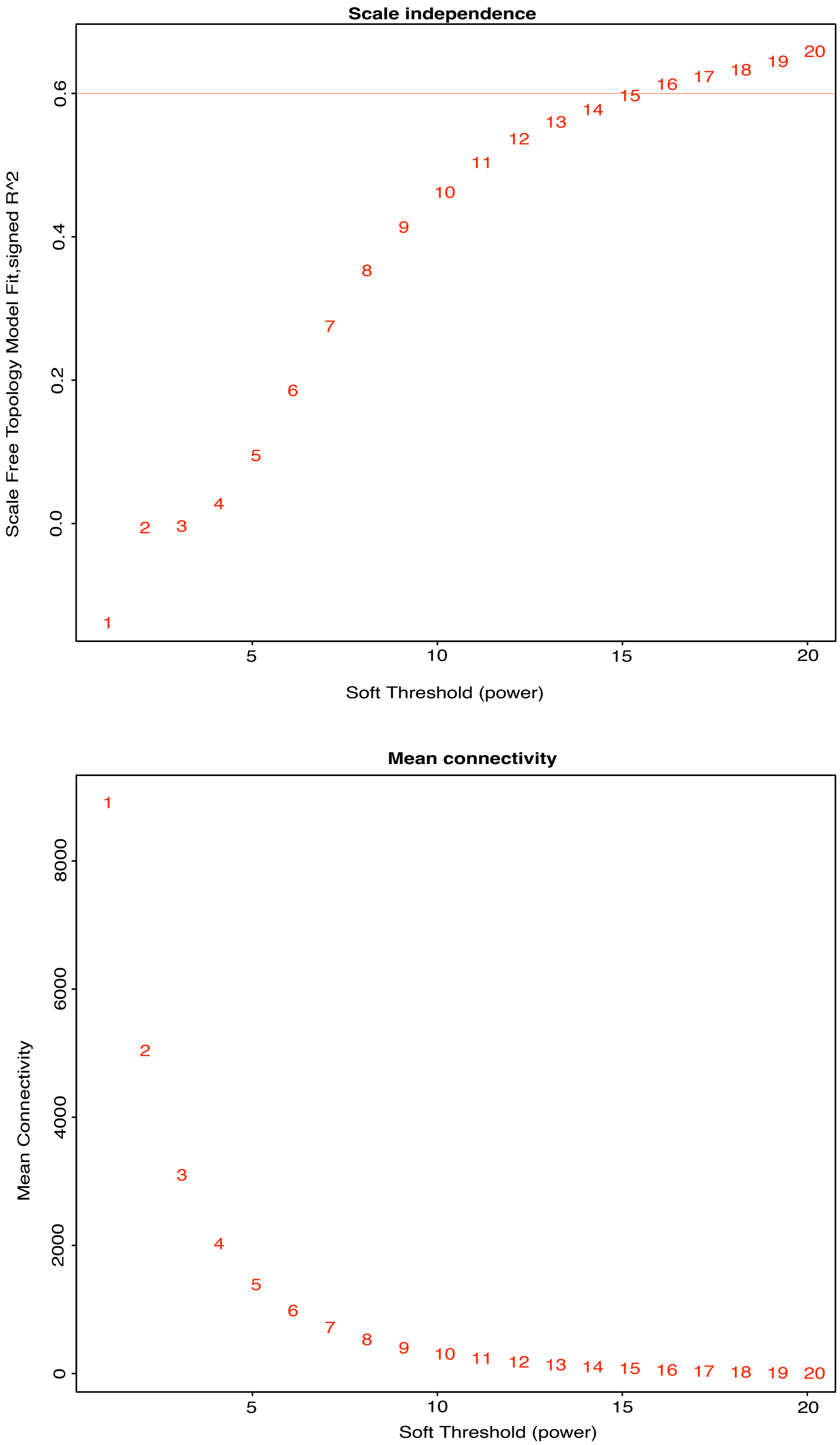


**Supplementary Figure S11: Picking a soft threshold for WGCNA based on scale independence and Mean connectivity.** Top: Scale-free topology index versus the power values 1 to 20; bottom: the mean connectivity versus the power values 1 to 20.

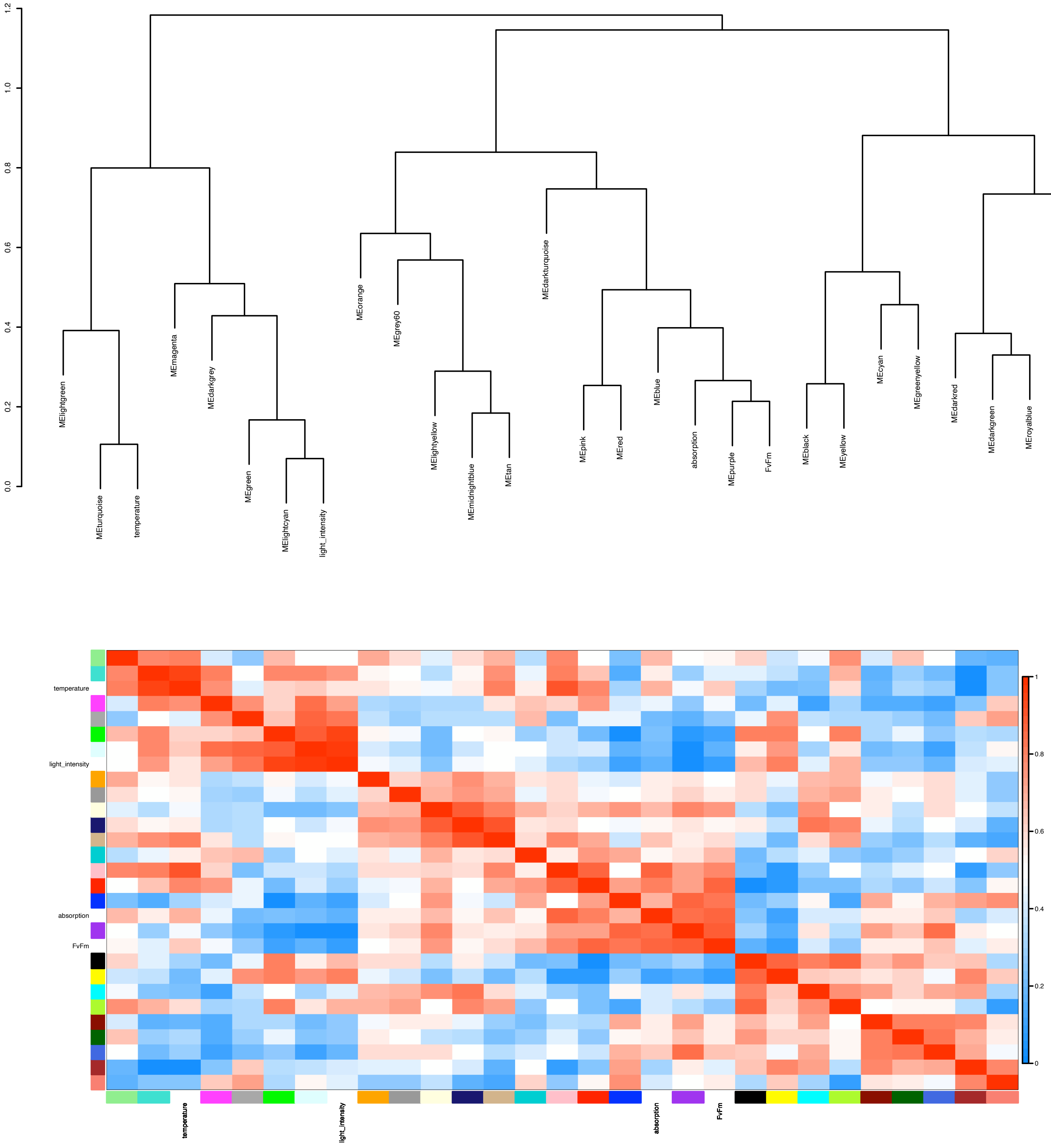


**Supplementary Figure S12: Clustering of different modules and traits.** Correlations between the modules (defined based on eigengenes) and the traits plotted as a heatmap from maximum correlation (1, red) to minimum (0, blue). The dendogram on the top shows the relationship between all 26 modules and the traits. The figure illustrates that a proper merging threshold was chosen. If the merging threshold is too small, many more modules would emerge, if overmerging were performed, unrelated genes would end up in one module.

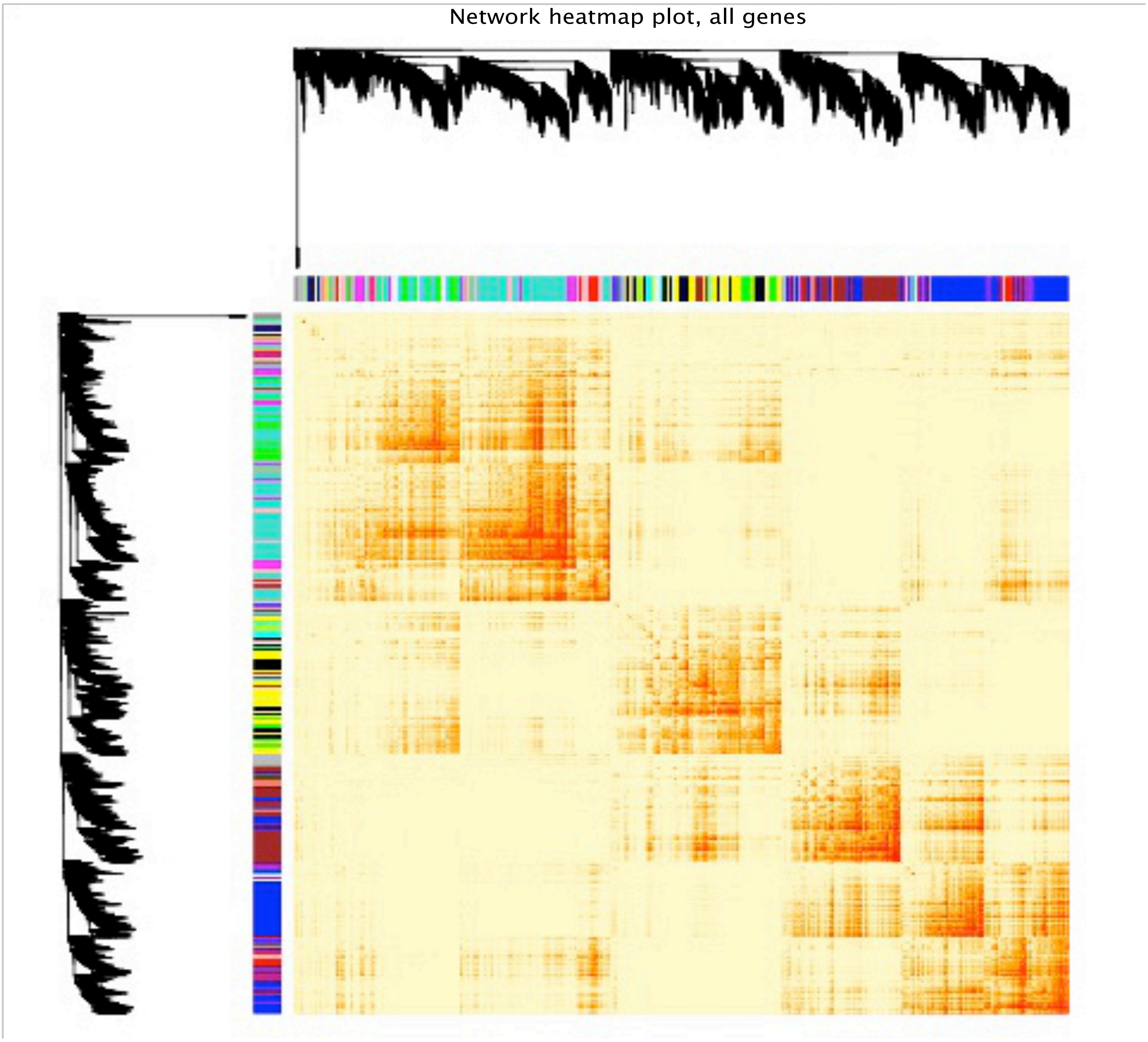


**Supplementary Figure S13: The graphical representation of the topological overlap matrix.** A plot of the connections of all genes versus all genes of Mesotaenium in the network. Dark orange represents

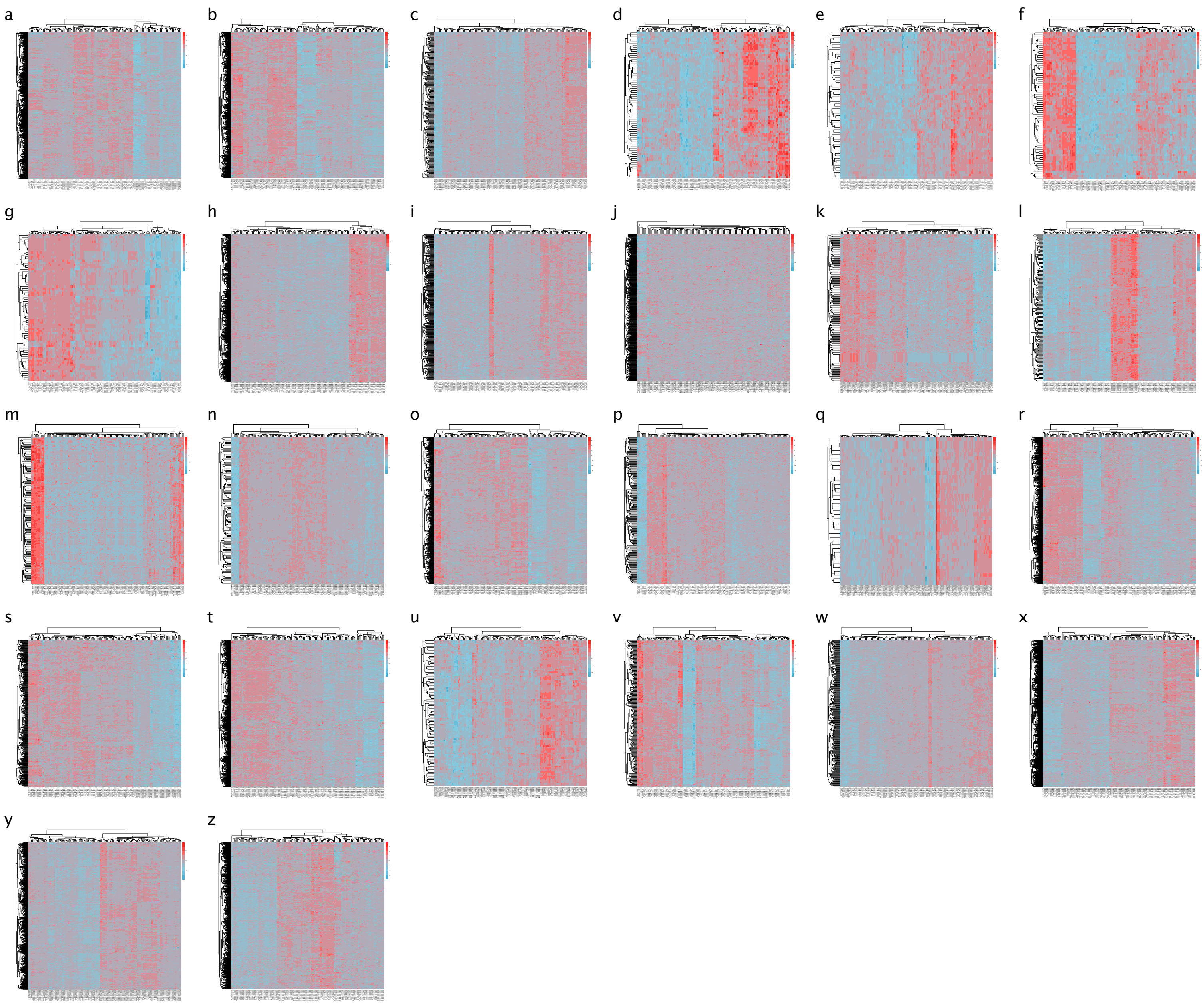


**Supplementary Figure S14: Heatmap of gene expression Z-score values for each module. (a)** blue **(b)** brown **(c)** cyan **(d)** darkgreen **(e)** darkgray **(f)** darkred **(g)** darkturquoise **(h)** green **(i)** green yellow **(j)** grey **(k)** grey60 **(l)** light cyan **(m)** light green **(n)** light yellow **(o)** magenta **(p)** midnight blue **(q)** orange **(r)** pink **(s)** purple **(t)** red **(u)** royal blue **(v)** salmon **(w)** tan **(x)** turquoise **(y)** yellow **(z)** black

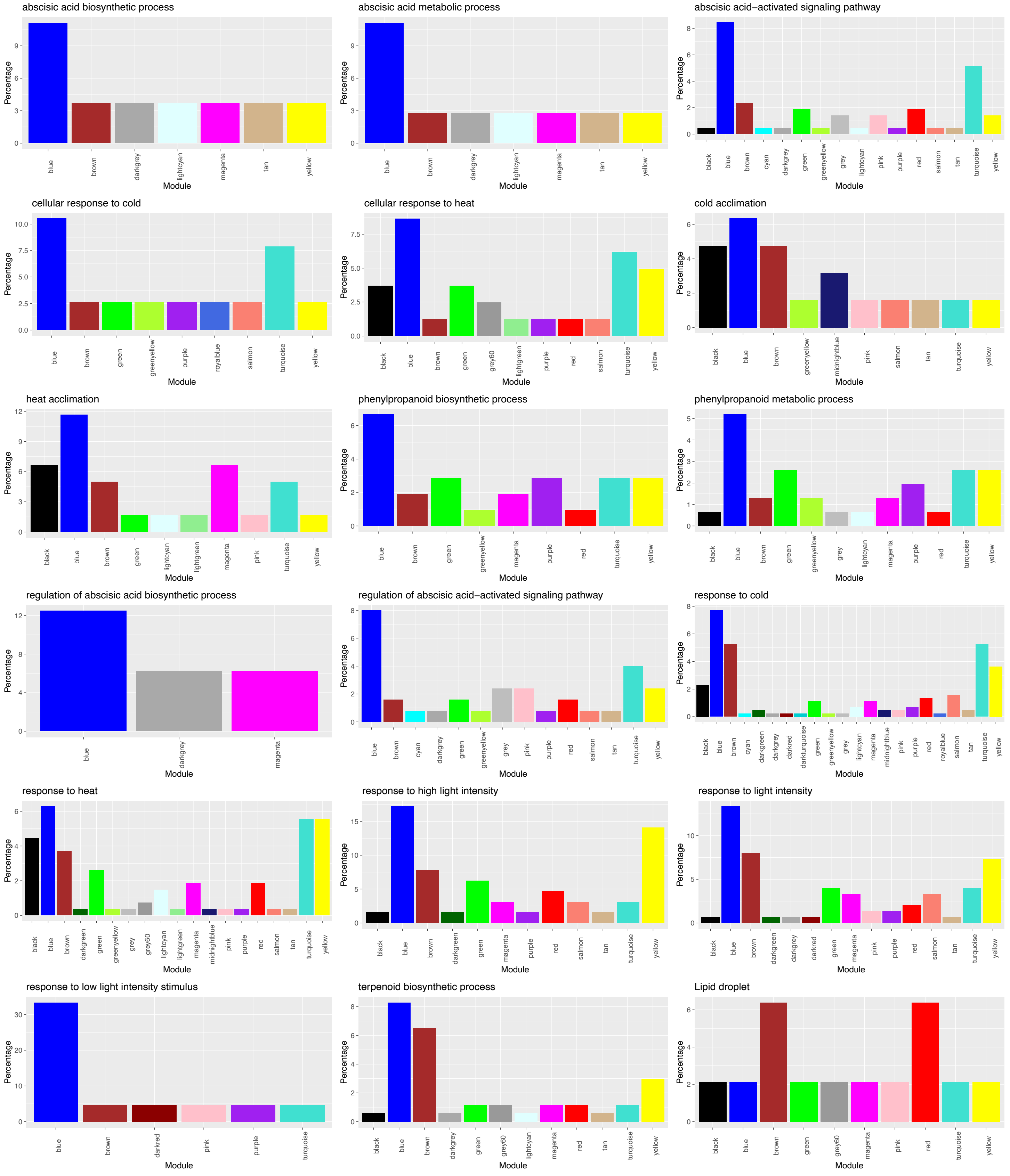


**Supplementary Figure S15: Distribution of of *A. thaliana* homologs for stress response genes among WGCNA modules.** Colors correspond to the 26 modules defined in main Figure 4a. Arabidopsis genes that were used as a query for homolog detection were based on keyword searches on TAIR and, in case of the lipid droplet-relevant genes, manually curated.

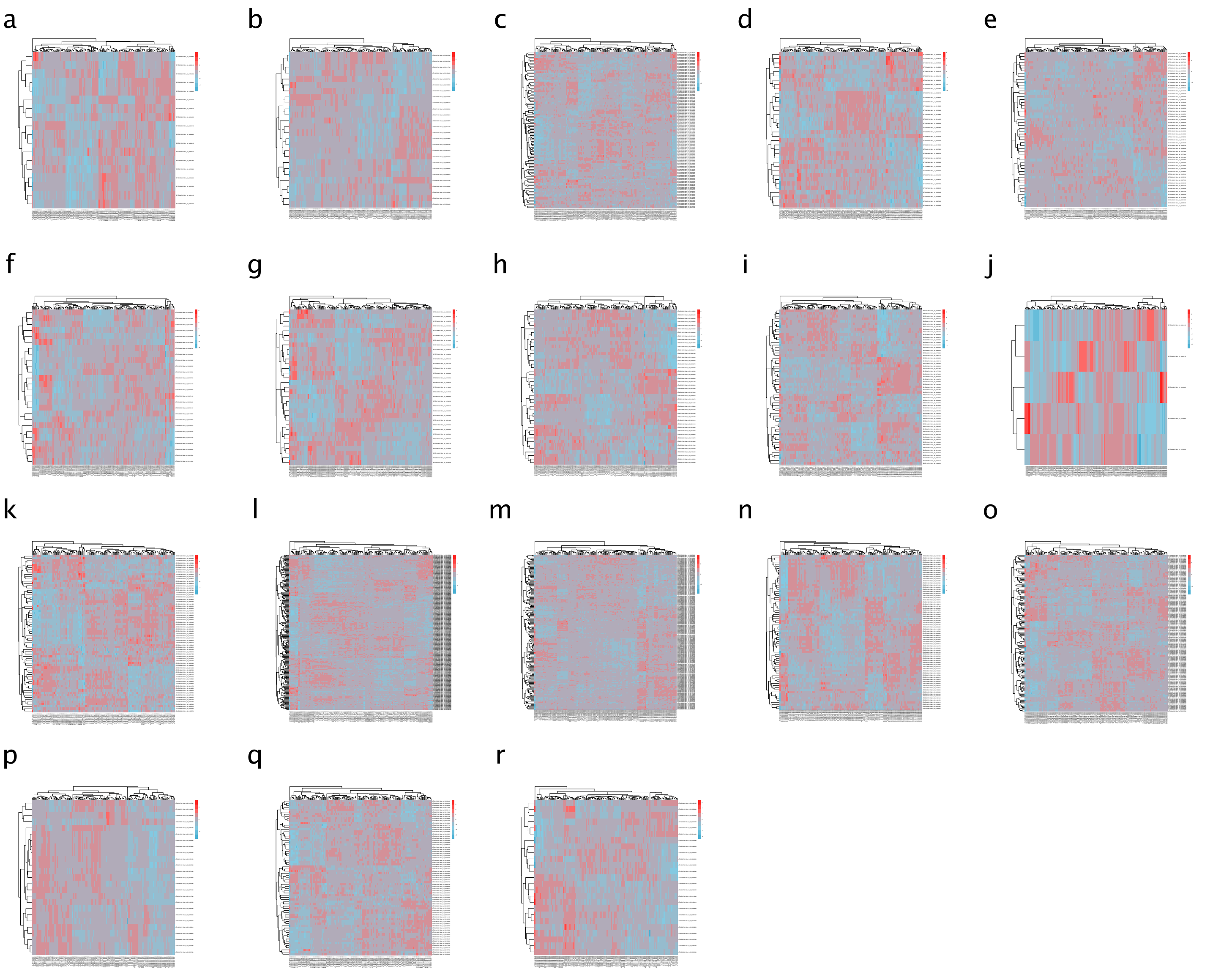


**Supplementary Figure S16: Heatmap of best blast hit of *A. thaliana* stress response genes in *M. endlicherianum* across different growth conditions.** **(a)** Abscisic acid biosynthetic process genes best blast hit gene expression **(b)** Abscisic acid metabolic process genes best blast hit gene expression **(c)** Abscisic acid−activated signaling pathway genes best blast hit gene expression **(d)** Cellular response to cold genes best blast hit gene expression **(e)** Cellular response to heat genes best blast hit gene expression **(f)** Cold acclimation genes best blast hit gene expression **(g)** Heat acclimation genes best blast hit gene expression **(h)** Phenylpropanoid biosynthetic process genes best blast hit gene expression **(i)** Phenylpropanoid metabolic process genes best blast hit gene expression **(j)** Regulation of abscisic acid biosynthetic process genes best blast hit gene expression **(k)** Regulation of abscisic acid−activated signaling pathway genes best blast hit gene expression **(l)** Response to cold genes best blast hit gene expression **(m)** Response to heat genes best blast hit gene expression **(n)** Response to high light intensity genes best blast hit gene expression **(o)** Response to light intensity genes best blast hit gene expression **(p)** Response to low light intensity stimulus genes best BLAST hit gene expression **(q)** Terpenoid biosynthetic process genes best blast hit gene expression **(r)** lipid droplet genes best blast hit gene expression

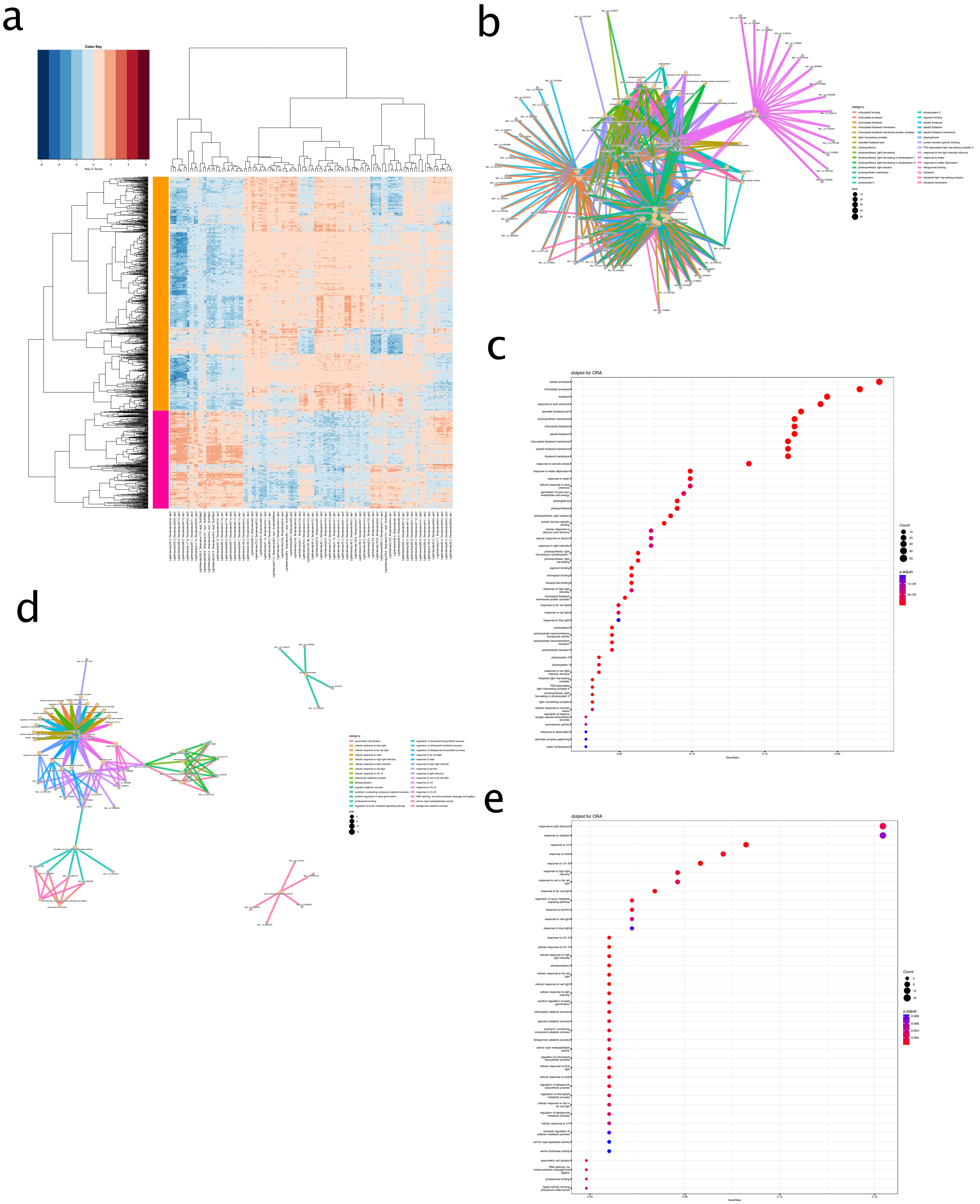


**Supplementary Figure S17: DEGs and GO-enrichment of Fv /Fm vs control. (a)** Heatmap of DEGs **(b)** cnetplot of orange cluster **(c)** dot plot of orange cluster **(d)** cnetplot of pink cluster **(e)** dot plot of pink cluster

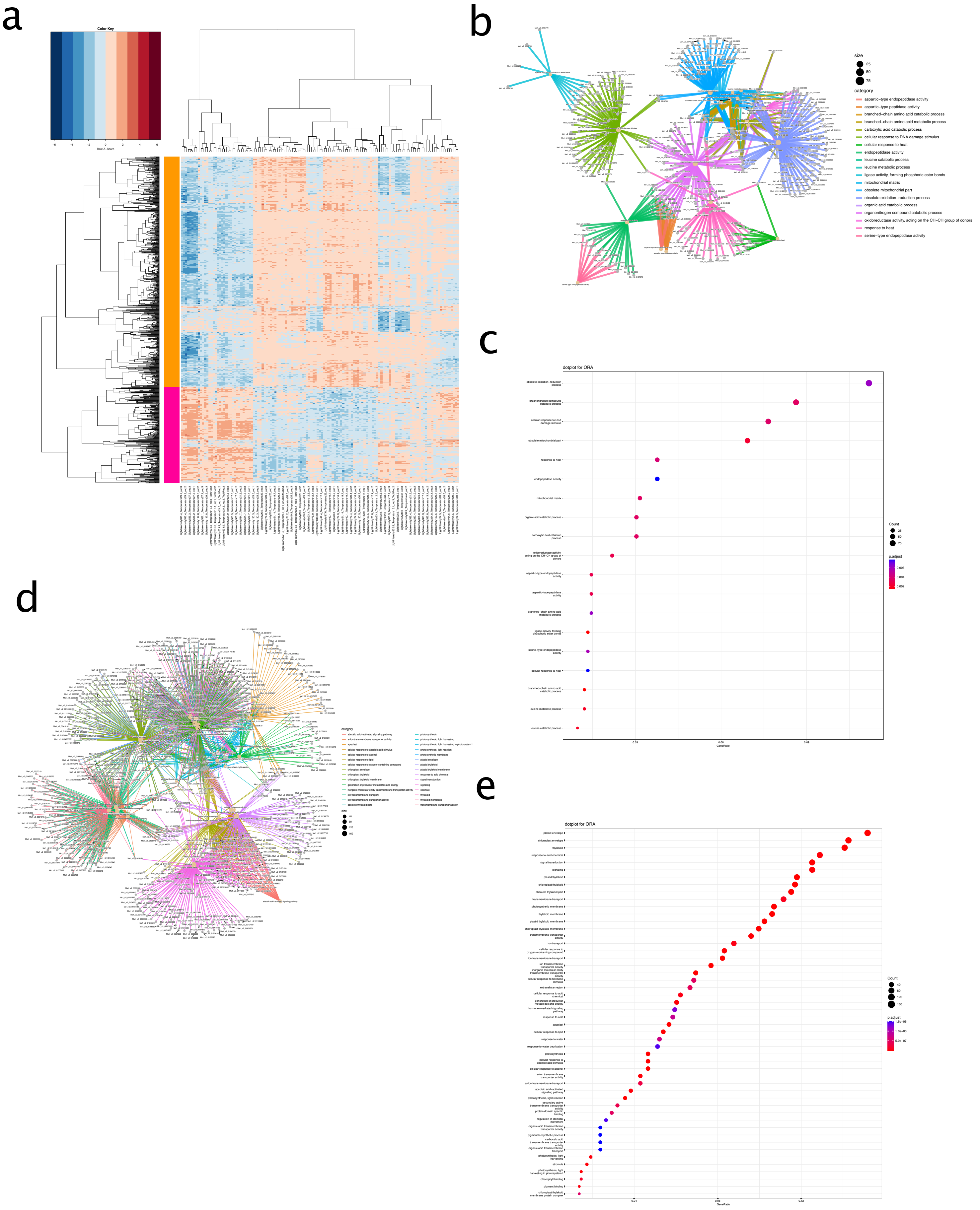


**Supplementary Figure S18: DEGs and GO-enrichment of HLI_HT vs. LLI_MT control. (a)** Heatmap of DEGs **(b)** cnetplot of orange cluster **(c)** dot plot of orange cluster **(d)** cnetplot of pink cluster **(e)** dot plot of pink cluster

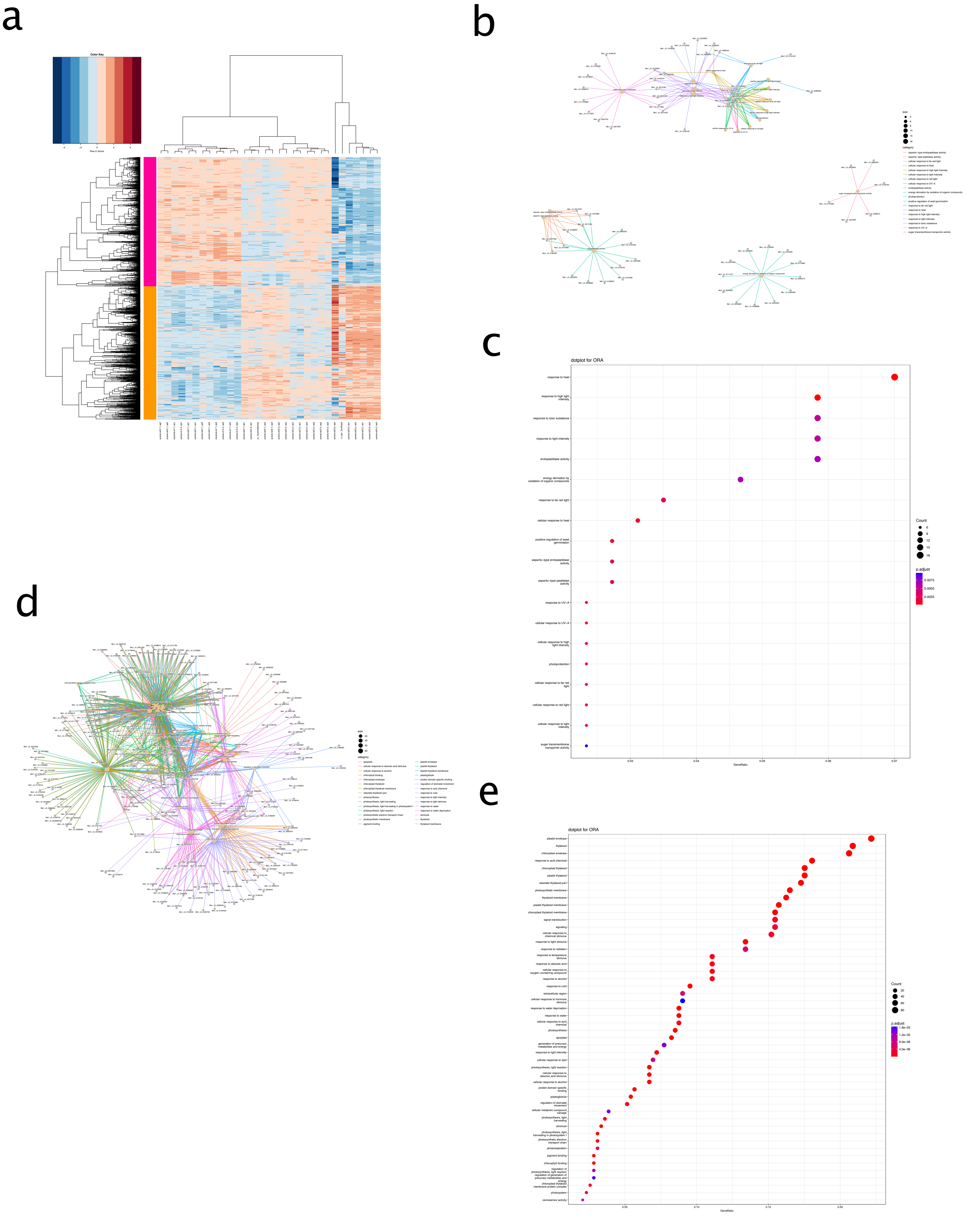


**Supplementary Figure S19: DEGs and GO-enrichment of MLI_HT vs. LLI_MT control. (a)** Heatmap of DEGs **(b)** cnetplot of orange cluster **(c)** dot plot of orange cluster **(d)** cnetplot of pink cluster **(e)** dot plot of pink cluster

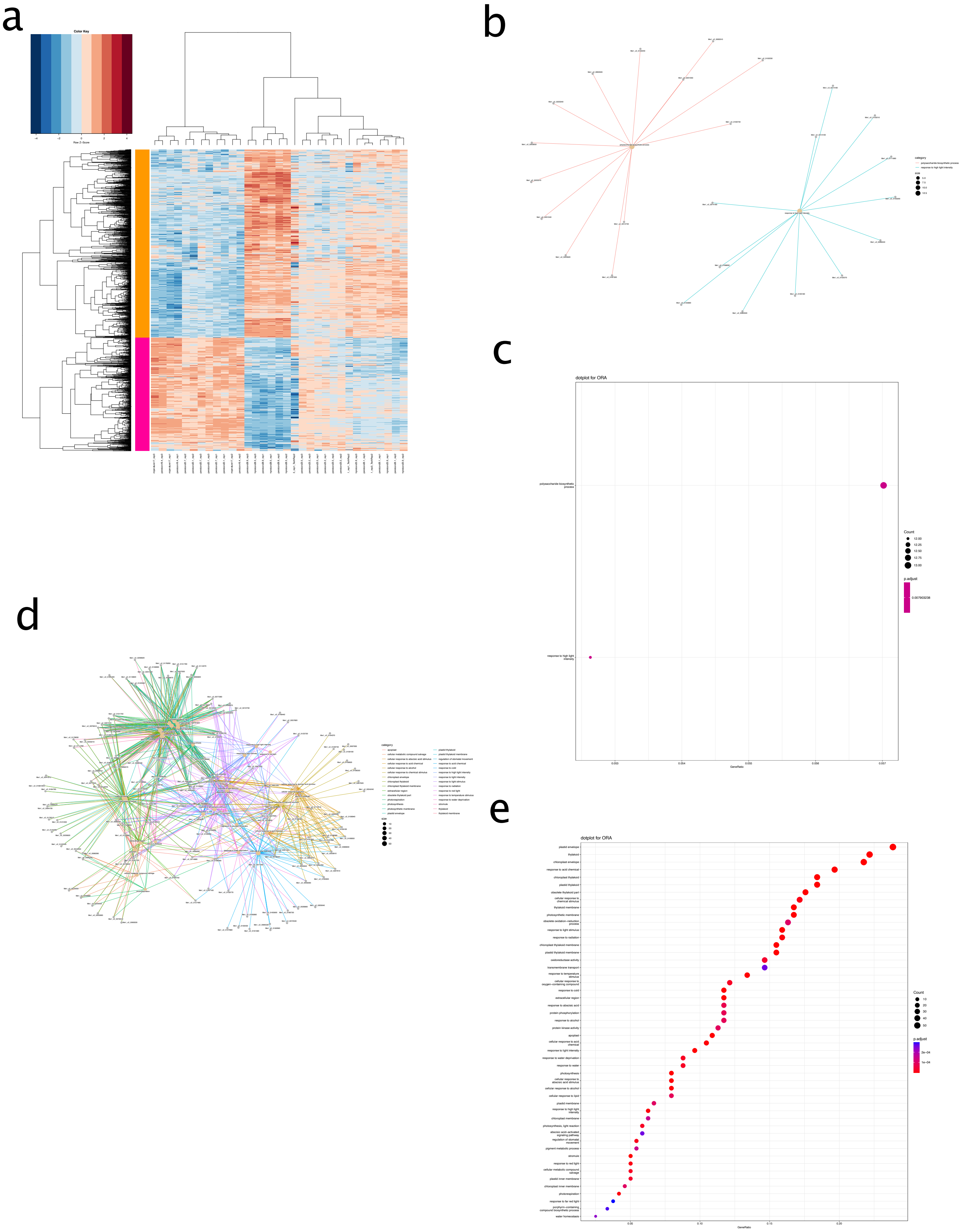


**Supplementary Figure S20: DEGs and GO-enrichment of LLI_HT vs. LLI_MT control. (a)** Heatmap of DEGs **(b)** cnetplot of orange cluster **(c)** dot plot of orange cluster **(d)** cnetplot of pink cluster **(e)** dot plot of pink cluster

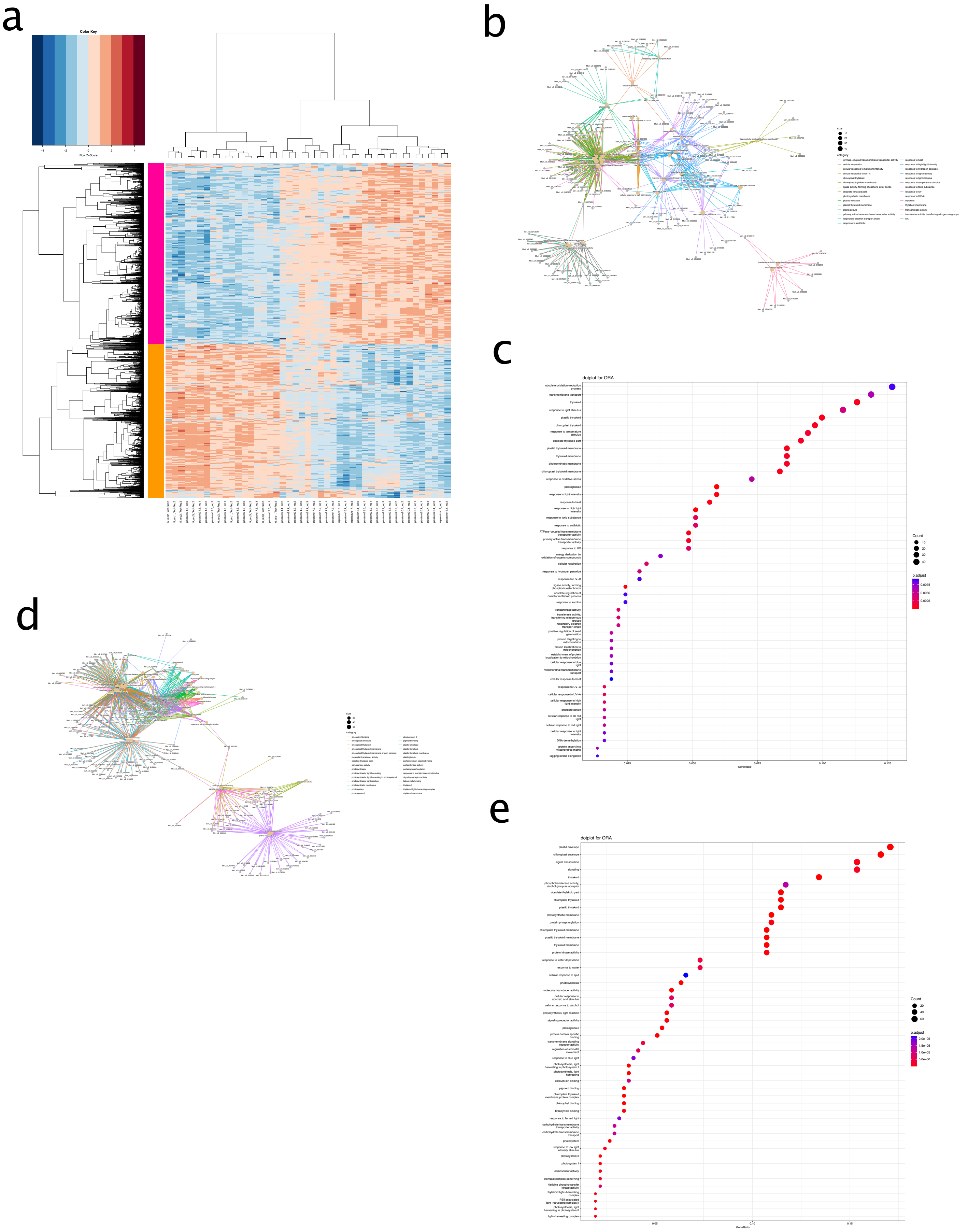


**Supplementary Figure S21: DEGs and GO-enrichment of HLI_MT vs. LLI_MT control. (a)** Heatmap of DEGs **(b)** cnetplot of orange cluster **(c)** dot plot of orange cluster **(d)** cnetplot of pink cluster **(e)** dot plot of pink cluster

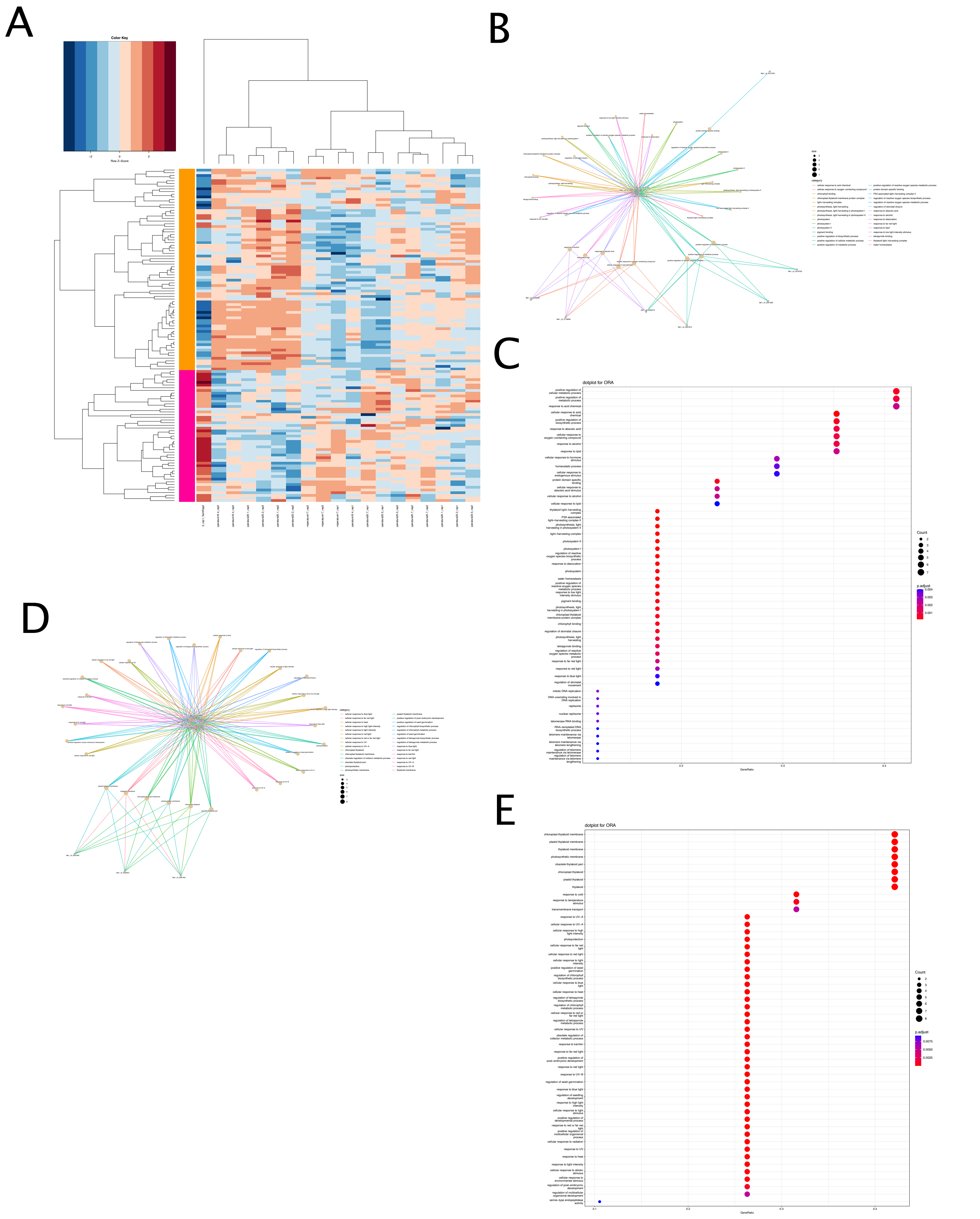


**Supplementary Figure S22: DEGs and GO-enrichment of MLI_MT vs. LLI_MT control. (a)** Heatmap of DEGs **(b)** cnetplot of orange cluster **(c)** dot plot of orange cluster **(d)** cnetplot of pink cluster **(e)** dot plot of pink cluster

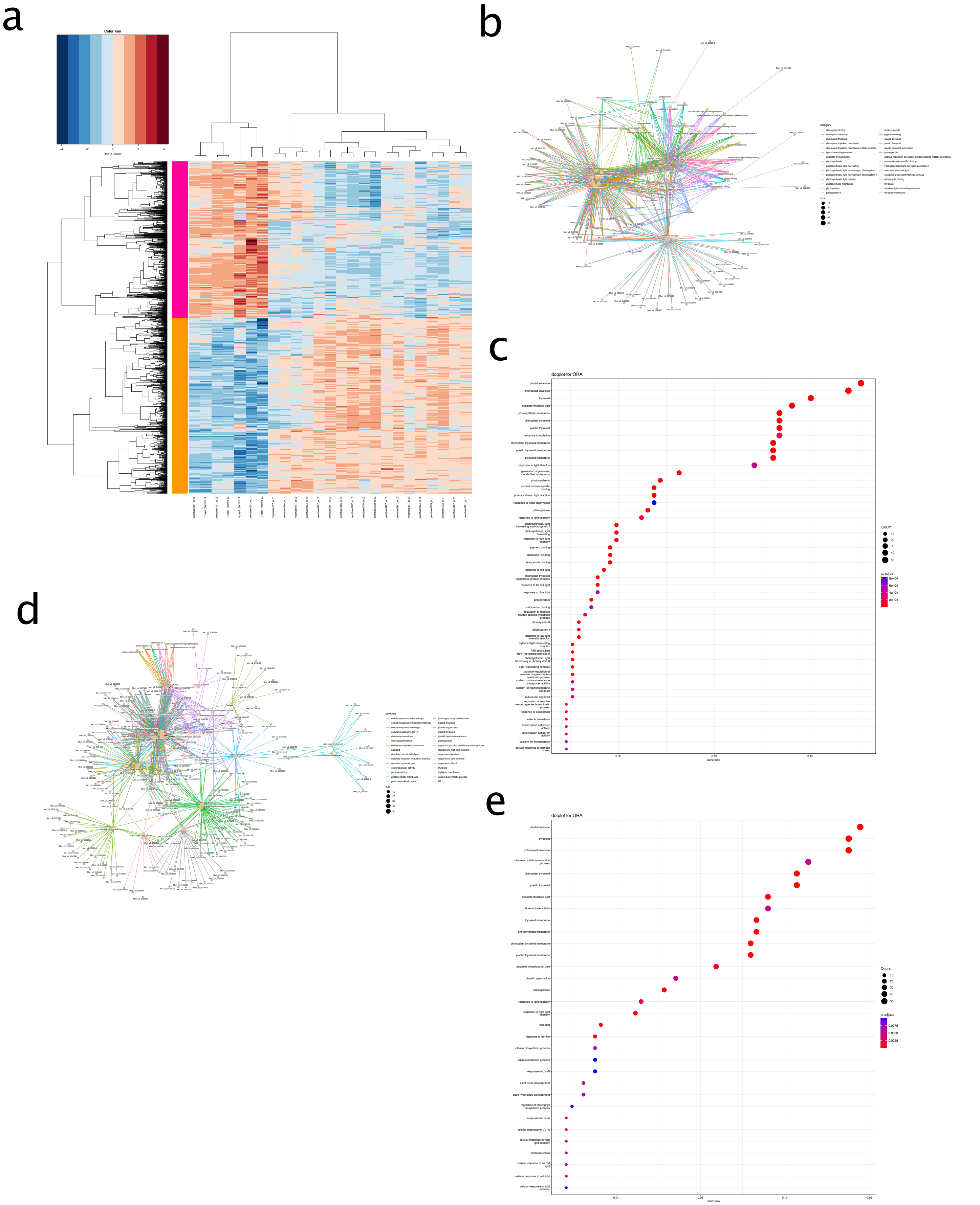


**Supplementary Figure S23: DEGs and GO-enrichment of HLI_LT vs. LLI_MT control. (a)** Heatmap of DEGs **(b)** cnetplot of orange cluster **(c)** dot plot of orange cluster **(d)** cnetplot of pink cluster **(e)** dot plot of pink cluster

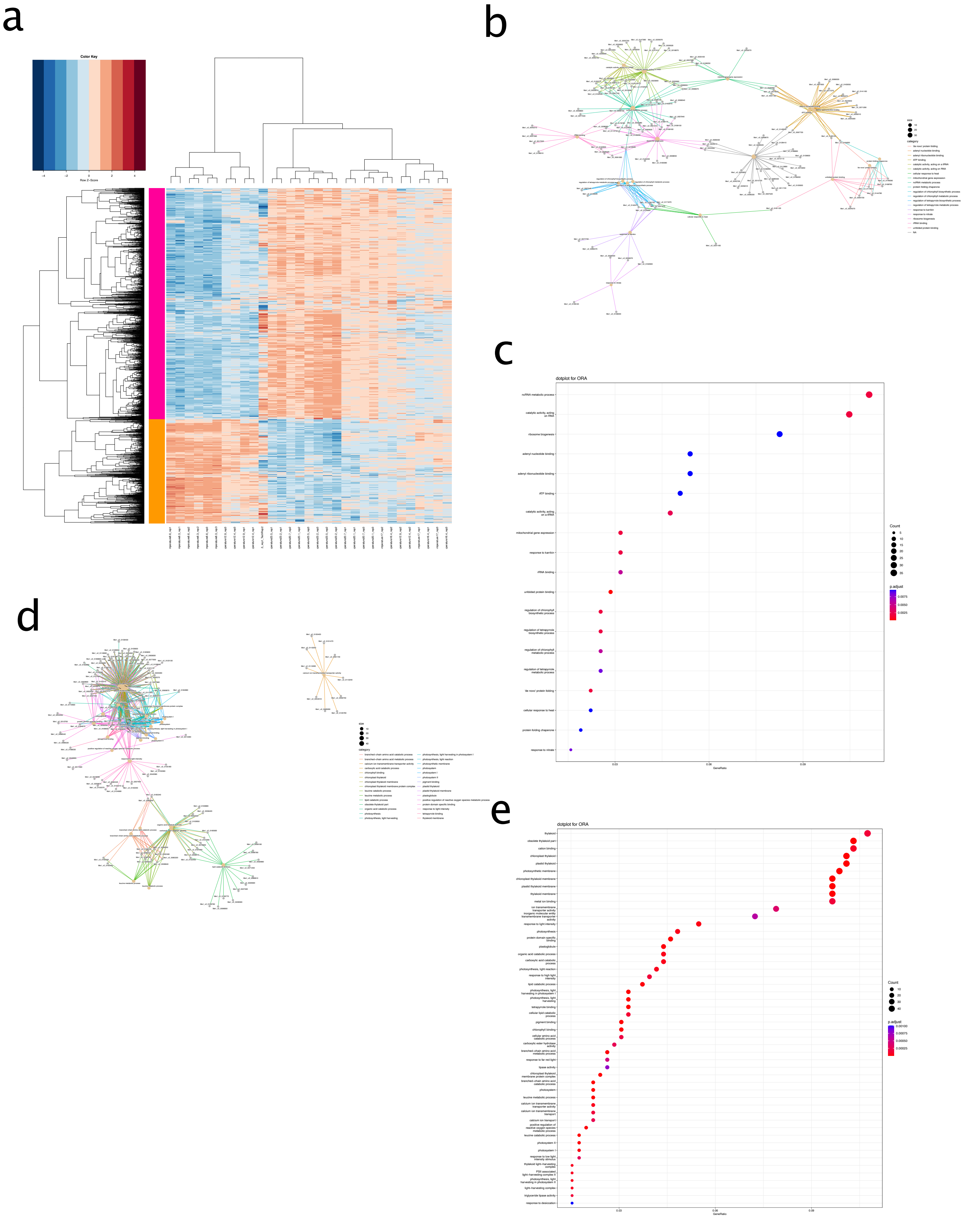


**Supplementary Figure S24: DEGs and GO-enrichment of MLI_LT vs. LLI_MT control. (a)** Heatmap of DEGs **(b)** cnetplot of orange cluster **(c)** dot plot of orange cluster **(d)** cnetplot of pink cluster **(e)** dot plot of pink cluster

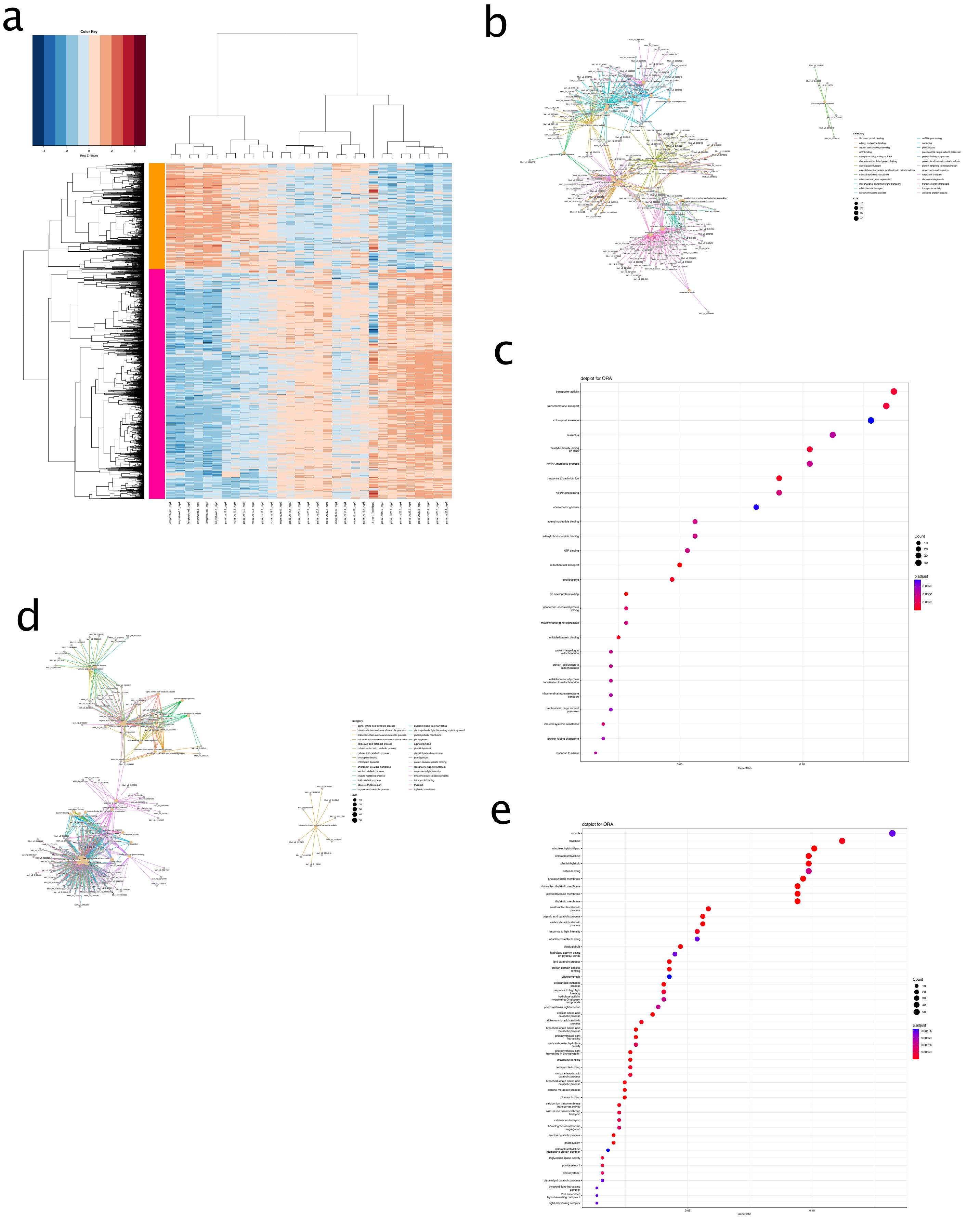


**Supplementary Figure S25: DEGs and GO-enrichment of LLI_LT vs. LLI_MT control. (a)** Heatmap of DEGs **(b)** cnetplot of orange cluster **(c)** dot plot of orange cluster **(d)** cnetplot of pink cluster **(e)** dot plot of pink cluster

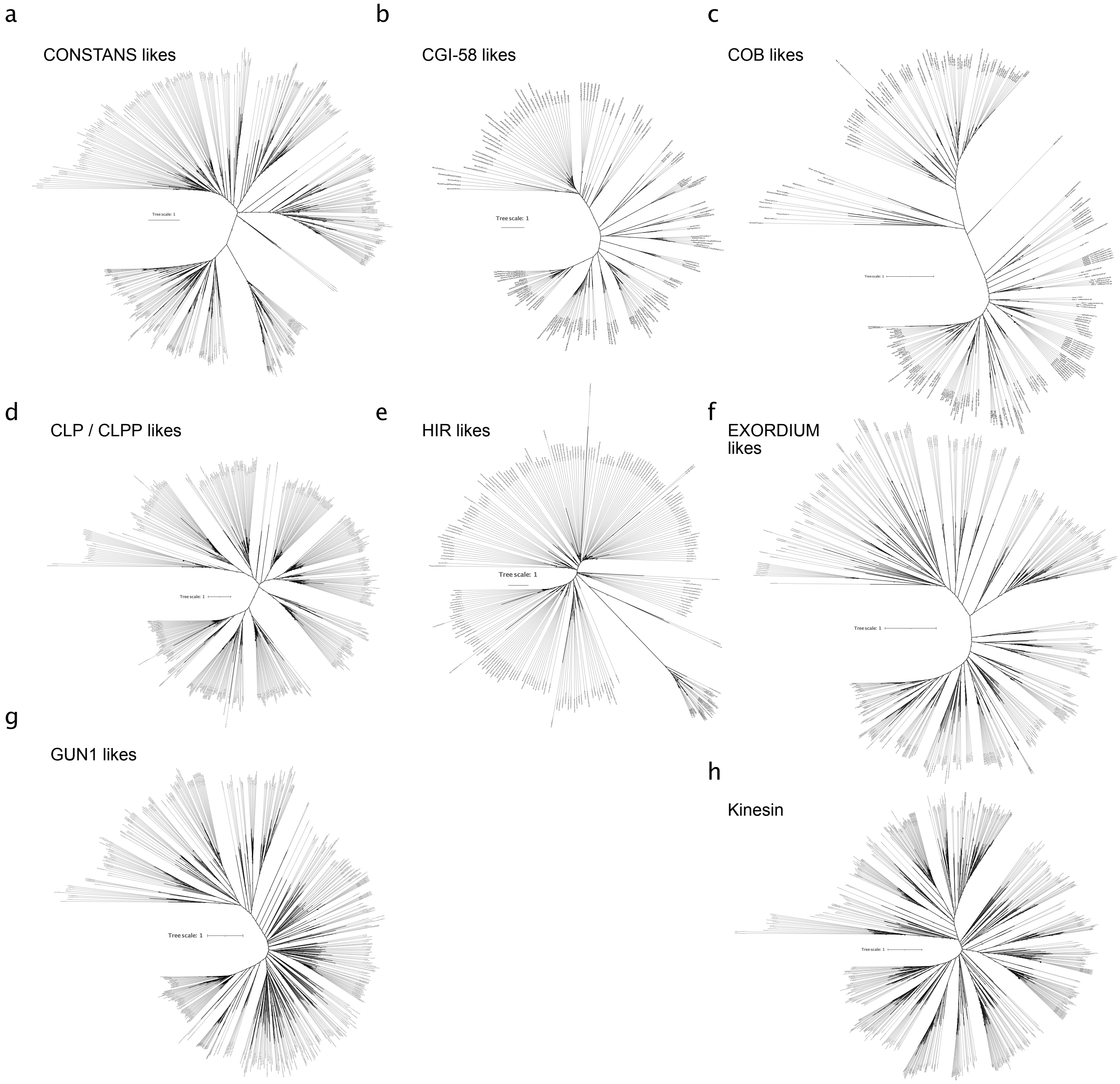


**Supplementary Figure S26: Fully-labeled phylogenies of hub genes.** **(a)** CONSTANS like **(b)** CGI-58 likes **(c)** COB likes **(d)** CLP / CLPP likes **(e)** HIR likes **(f)** EXORDIUM likes **(g)** GUN1 likes **(h)** Kinesin. All phylogenies were computed with IQ-TREE multicore version 1.5.5, their respective best model according to Bayesian Information Criterion and 1000 ultrafast bootstrap replicates.

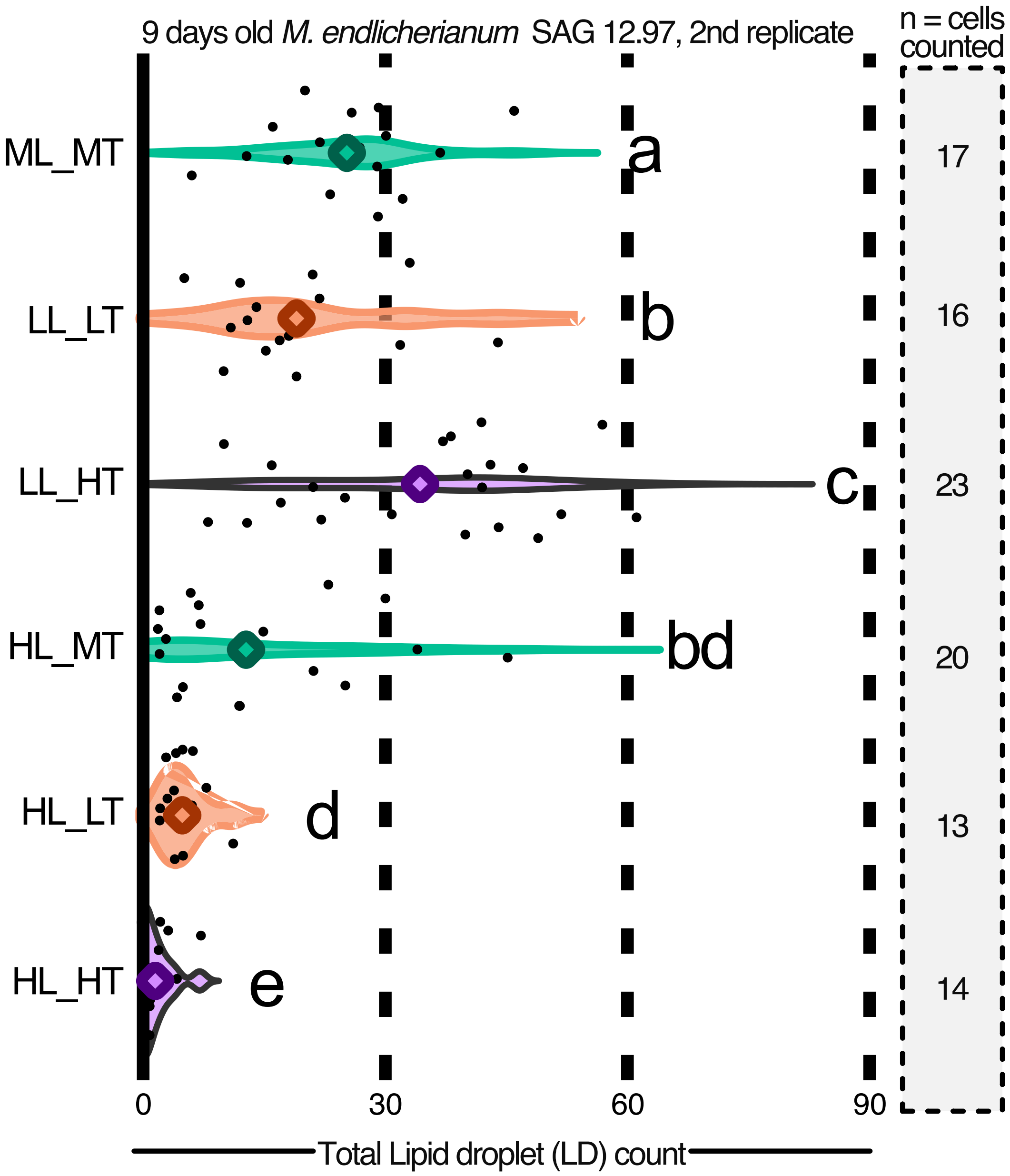


**Supplementary Figure S27: Lipid droplet count setup 2.** Violin plots of second biological replicate of LD quantification after 9 days of exposure to different environmental conditions including statistical analysis using Mann-Whitney U statistics (significance grouping based on p value < 0.05).

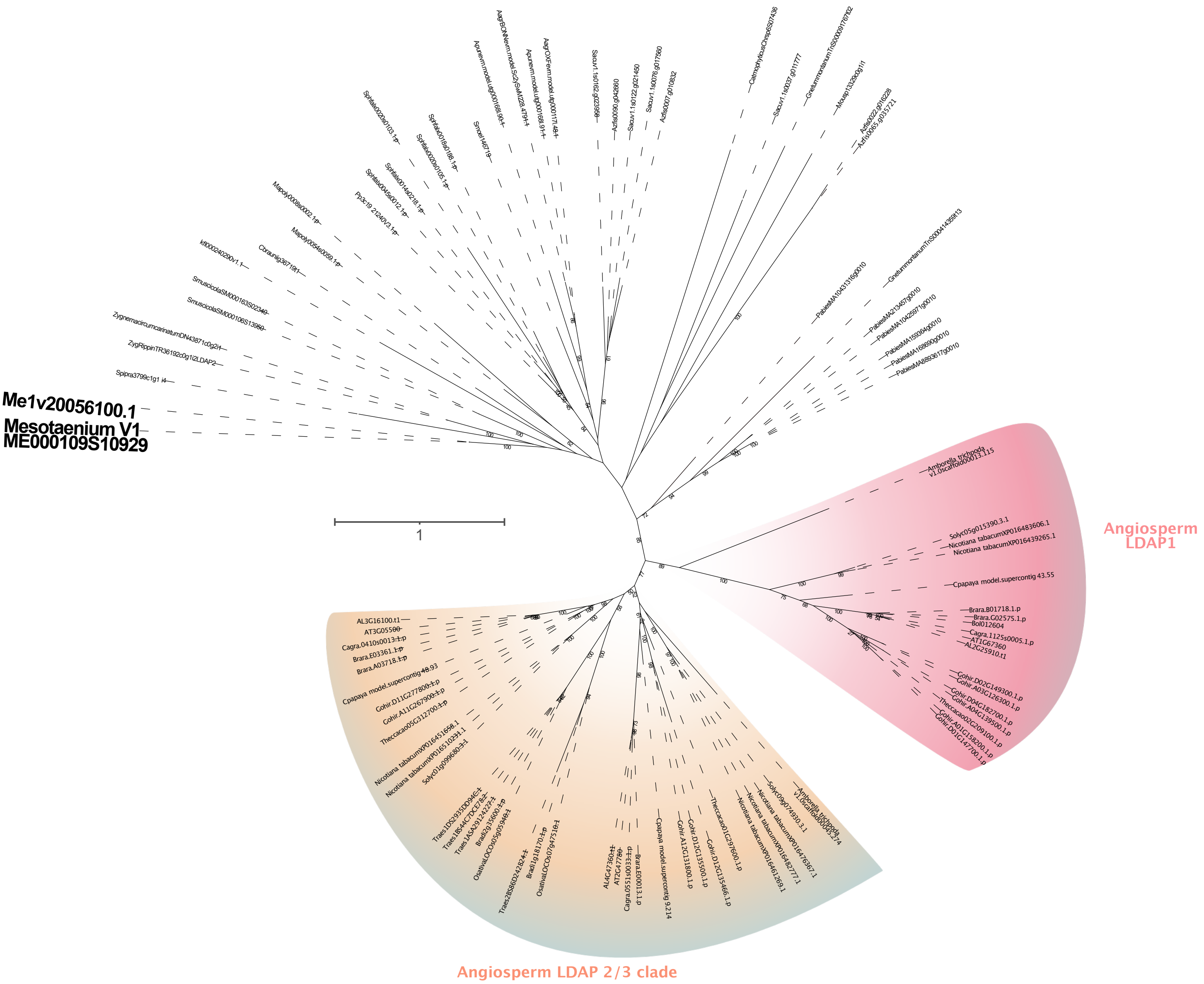


**Supplementary Figure S28: LDAP phylogeny.** Maximum likelihood phylogeny computed with IQ-TREE multicore version 1.5.5 based on a MAFFT L-INS-I alignment. Using ModelFinder, we determined JTT+I+G4 as best model for protein evolution based on Bayesian Information Criterion. The phylogeny was based on the alignment in de Vries and Ischebeck (2020) after adding the annotation V2 homolog of MeLDAP (Me1v20056100.1).
